# A new chemotype of chemically tractable nonsteroidal estrogens based on a thieno[2,3-*d*]pyrimidine core

**DOI:** 10.1101/2022.04.15.488344

**Authors:** Vamshikrishna Reddy Sammeta, John D. Norris, Sandeep Artham, Chad D. Torrice, Jovita Byemerwa, Carstyn Joiner, Sean W. Fanning, Donald P. McDonnell, Timothy M. Willson

## Abstract

Despite continued interest in development of nonsteroidal estrogens and antiestrogens, there are only a few chemotypes of estrogen receptor ligands. Using targeted screening in a ligand sensing assay we identified a phenolic thieno[2,3-*d*]pyrimidine with affinity for estrogen receptor α. An efficient three-step synthesis of the heterocyclic core and structure-guided optimization of the substituents resulted in a series of potent nonsteroidal estrogens. The chemical tractability of the thieno[2,3-*d*]pyrimidine chemotype will support the design of new estrogen receptor ligands as therapeutic hormones and antihormones.

**GRAPHICAL ABSTRACT:** 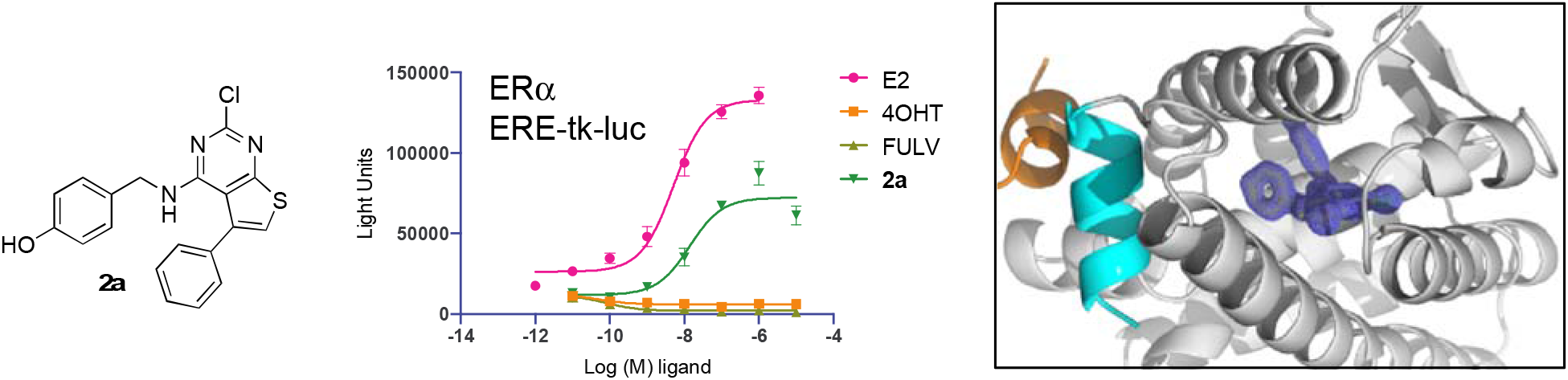

## INTRODUCTION

Estrogens are important endocrine sex hormones that control many aspects of female physiology and have important actions in males.^1^ Estrogen pharmacology is primarily mediated through the estrogen receptor α (ERα), a ligand-activated transcription factor and member of the nuclear receptor superfamily.^2-3^ A second estrogen receptor β (ERβ), which shares high sequence identity with ERα, has also been characterized. However, unlike ERα, the contribution of ERβ to the pharmacology of estrogens remains contentious despite extensive study over a 20-year period.^4^ ERα functions as a homodimer to bind to palindromic DNA sequences, known as estrogen responsive elements (ERE), in the enhancer region of its target genes. The binding of an estrogen agonist (hormone) to the ligand binding domain (LBD) of the ERα homodimer facilitates the recruitment of coactivators to the receptor-DNA complex and initiation of gene transcription. ERβ can form heterodimers with ERα and one of its functions may be to modulate the breadth of target gene activation in tissues where the receptors are coexpressed.^5-6^

The main circulating estrogens in women are estrone (E1) and estradiol (E2), although additional metabolites with lower activity have been detected.^7^ All of the naturally-occurring steroidal estrogens have an aromatic A-ring with a phenol at C-3 (Figure 1). Chemical modification of the steroid core at C-7 with a long lipophilic chain results in compounds with antagonist (antihormone) activity, such as the breast cancer drug fulvestrant (FULV), which blocks the action of endogenous estrogens and has the additional effect of promoting receptor turnover in cells.^8^ Diethylstilbestrol (DES) is a synthetic nonsteroidal estrogen that was first reported in 1938.^9^ DES contains two phenols to mimic the steroid A- and D-rings and its affinity for ERα is similar to E2. Many nonsteroidal antiestrogens have been developed from the DES core, including 4-hydroxytamoxifen (4OHT) where addition of a third aromatic substituent was used to modulate receptor activation.^10^ Several cyclic analogs of 4OHT that block estrogen action or induce ERα degradation remain in clinical development as breast cancer drugs.^11^ A wide range of phenol-containing natural products have also been shown to possess estrogenic activity,^12^ including the phytoestrogen ferutinin and the mycotoxin β-zearalenol. In addition, several phenolic plasticizers and detergents, such as butylate hydroxy anisole (BHA), bisphenol A (BPA), and nonylphenol have weak estrogenic activity and are labelled as potential xenoestrogens due their occurrence in the environment.^13^ It is notable that most synthetic estrogens and antiestrogens contain a phenol as a common pharmacophore (Figure 1).^10, 14^ Remarkably, despite decades of pharmaceutical research on synthetic ERα ligands, there is a dearth of chemically tractable chemotypes. Most nonsteroidal ligands are still based on the DES/4OHT core with only limited chemical diversity in their structures.^10, 14^ Given the continued interest in development of nonsteroidal ERα ligands for hormone replacement or breast cancer therapy we sought to identify new chemotypes of synthetic estrogens.

**Figure 1.**
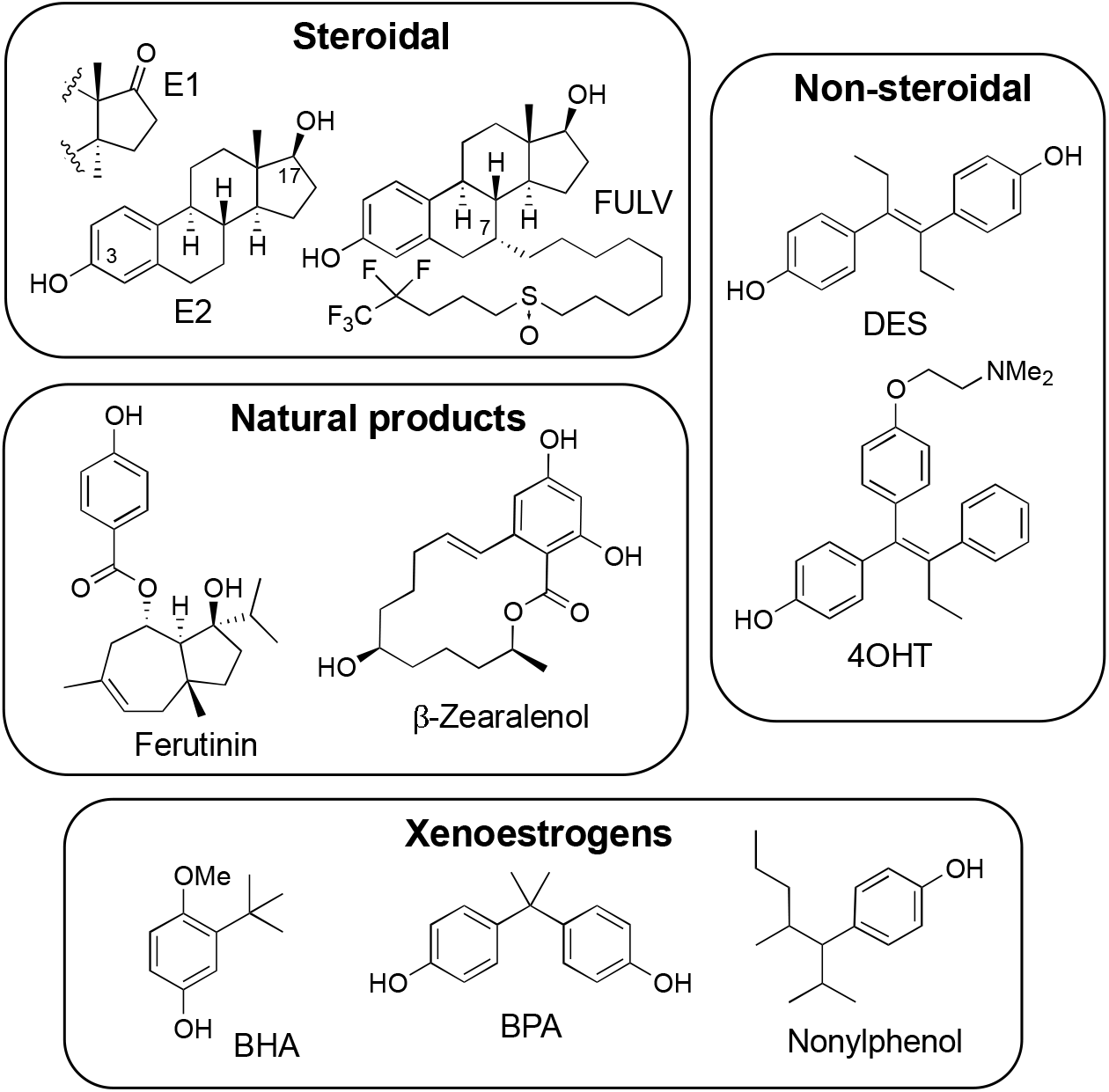
Chemotypes of estrogen receptor ligands: E1, estrone. E2, 17β-estradiol. FULV, fulvestrant. DES, diethylstilbestrol. 4OHT, 4-hydroxytamoxifen. BHA, butylated hydroxyanisole. BPA, bisphenol A. The phenol pharmacophore is highlighted in yellow and red in each compound.

## RESULTS AND DISCUSSION

### Identification of a thieno[2,3-*d*]pyrimidine ERα agonist

To identify new cell active ERα ligands, we developed a sensitive, high throughput screening assay in 384-well format. We had previously identified an 11 amino acid peptide (αII, SSLTSRDFGSWYASR) that was recruited to ERα when bound by small molecule agonists, partial agonists, and antagonists.^15-16^ To configure the assay in a two-hybrid format, HepG2 cells were transiently transfected with expression vectors for ERα-VP16 and αII-Gal4 fusions as well as a luciferase reporter construct under the control of five copies of a Gal4 upstream enhancer element. Addition of a wide range of standard hormones and antihormones resulted in an increase in luciferase (Figure 2A), demonstrating that the assay was sensitive to ER ligands independent of their agonist or antagonist functional activity. The ERα ligand sensing assay (ERα LiSA) demonstrated a robust Z’ = 0.7 in a 384-well format that was suitable for high throughput screening.

**Figure 2.**
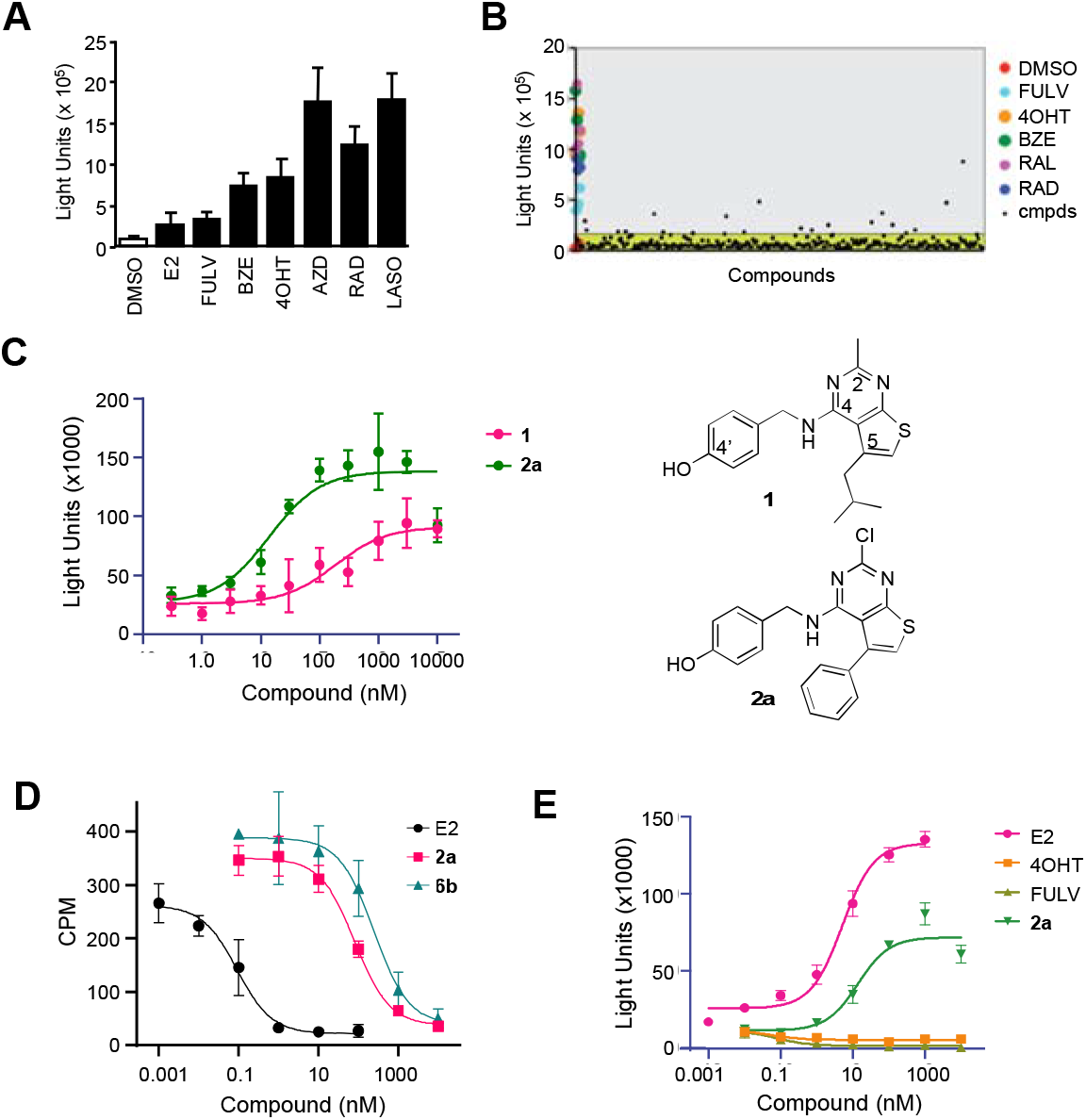
**A**. ERα LiSA in HepG2 cells transfected with Gal4-αII, ERα-VP16, and 5x-gal-tata-Luc. After 24h, cells were lysed and luciferase assays were performed. **B**. ERα LiSA screen. Representative data from plate 1/28. Compounds with luciferase >4x (grey shading) over background (green shading) were selected as primary hits. **C**. Dose-response of the thieno[2,3-*d*]pyrimidine hits **1** and analog **2a** in the ERα LiSA. **D**. Whole cell ERα affinity determined by competition binding with ^3^H-E2. **E**. Agonist activity in HepG2 cells transfected with ERα, 7x-TK-ERE-Luc, and Renilla luciferase. All data: error bars represent ± SD of triplicate points. BZE, bazedoxifene; AZD, AZD-9496; RAD, RAD-1901; LASO, lasofoxifene.

**Figure 4.**
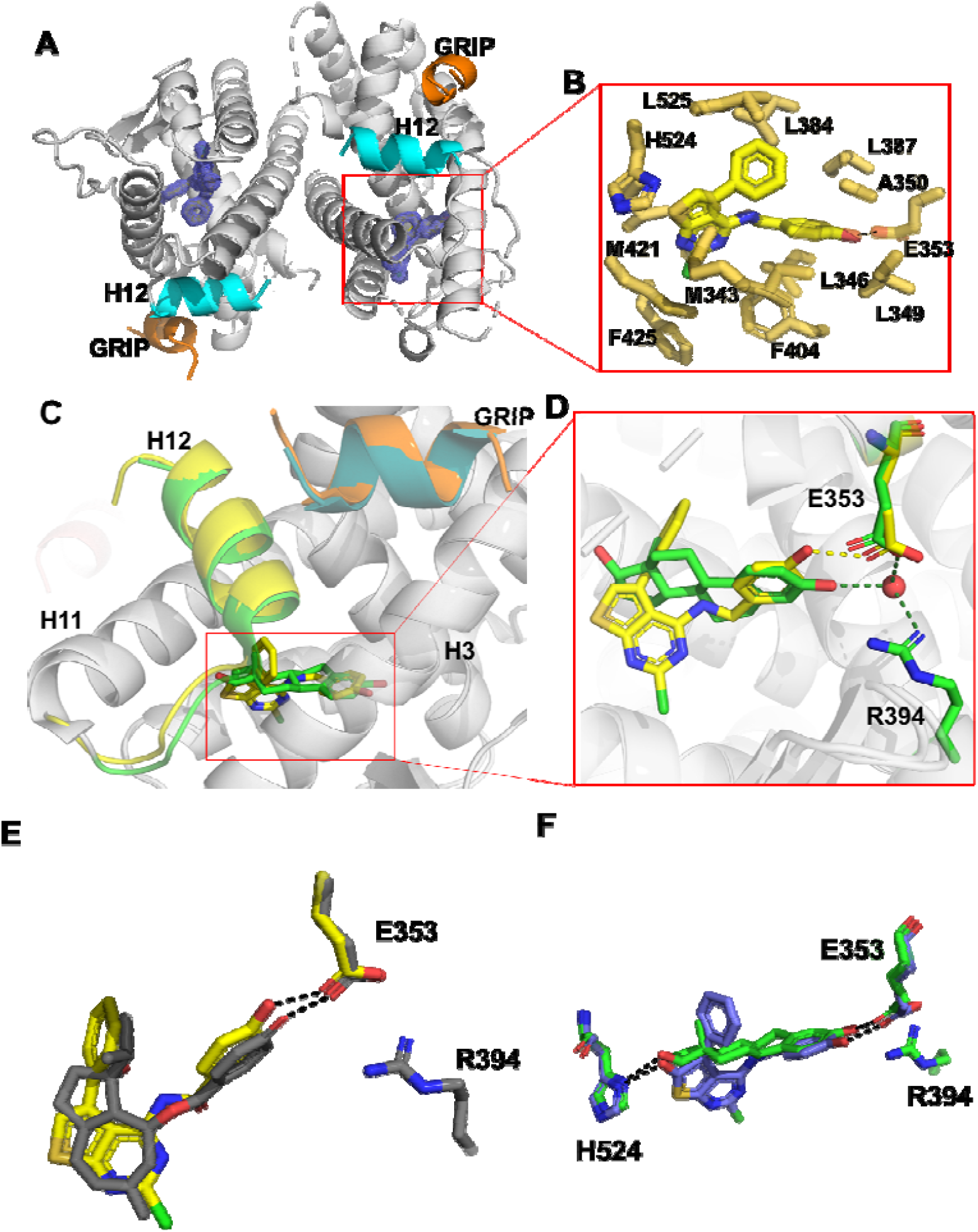
Structural Basis of ERα agonist activity of thieno[2,3-*d*]pyrimidines **2a** and **6b. A**. Overview of the co-crystal structure with ERα LBD with 2mFo-DFc difference maps of **2a** in the ligand binding pocket contoured to 2σ. **B**. Thieno[2,3-*d*]pyrimidine binding pocket. Amino acid sidechains within 4Å of **2a** shown in bold. **C**. Superposition with E2 (green) and **2a** (yellow) highlighting the differences between ligand binding poses. **D**. Expanded view of the differences in phenolic hydrogen bonding patterns between **2a** and E2. **E**. Superposition of ferutinin (grey) and **2a** (yellow) showing the similar binding poses. **F**. Superposition of E2 (green) and **6b** (purple) highlighting the H-bond interactions with E353 and H524. All highlighted protein features are color coded to their respective ligands. Dashed lines represent hydrogen bonds. The PDB codes are 6CBZ and 4MG7 for the E2 and ferutinin structures, respectively.

We opted to perform a targeted diversity screen, based on the principal that natural and synthetic ER ligands often contain a phenol in their core structure (Figure 1). A library 8400 phenols arrayed across 28 384-well plates was obtained from Enamine (Monmouth Jct., NJ) representing a subset of their >2.9 million compound screening collection. All compounds were screened at 10 μM final concentration in the ERα LiSA and results presented as a scatter plot (Figure 2B). Screening the full library of 8400 compounds yielded 324 primary hits that were confirmed in follow-up 4-point dose response studies (SI Table 1). Blinded positive control compounds raloxifene (RAL, ERα antagonist) and diethylstilbestrol (DES, ERα agonist) scored as the top hits during the screening process validating our general approach (SI Table 1). The primary hits were triaged by potency to yield 34 compounds with EC_50_ <1.0 µM in the ERα LiSA. A thieno[2,3-*d*]pyrimidine **1** (Figure 2C) (EC_50_ = 180 nM) was selected for further optimization based on the novelty of the core structure and the commercial availability of 26 analogs from Chemspace (Monmouth Jct., NJ). While these additional analogs did not provide a systematic evaluation of the thieno[2,3-*d*]pyrimidine core, some initial structure-activity trends were observed (SI Table 2): switching the isobutyl group on C-5 on the thieno[2,3-*d*]pyrimidine core to a phenyl group resulted in a 2-fold increase in potency; the 4-hydroxybenzylamine at C-4 on the thienopyrimidine appeared to be essential for activity, since analogs where the 4’-OH was replaced with 3’-OH, 4’-NH_2_ or 4’-CO_2_H were >10-fold less active; and at C-2, chlorine was the only replacement for methyl that had improved activity. Thieno[2,3-*d*]pyrimidine (**2a**) was the most potent of these analogs with EC_50_ = 14 nM in the ERα LiSA (Figure 2C). To confirm that the activity of **2a** in the ERα LiSA was consistent with direct interaction on the receptor, a whole-cell competition binding assay was performed using ^3^H-E2 as a radioligand. Thienopyrimidine **2a** competed for ^3^H-E2 binding to ERα with an IC_50_ of 65 nM (Figure 2D). For comparison, unlabelled E2 competed with an IC_50_ = 0.2 nM.

The functional activity of **2a** was determined in an estrogen-responsive reporter gene assay using HepG2 cells that were transiently transfected with expression plasmids for ERα and an (ERE)_7_-tk-luciferase reporter. Thieno[2,3-*d*]pyrimidine (**2a**) induced a dose responsive increase in luciferase activity to 60-75% of the maximal efficacy of the natural hormone E2 (Figure 2E). The nanomolar agonist activity of thienopyrimidine **2a**, EC_50_ = 14 nM, was consistent with its affinity as an ERα ligand. The binding and functional data characterized thieno[2,3-*d*]pyrimidine (**2a**) as a new chemotype of nonsteroidal estrogens.

### Cocrystal structure of 2a with ERα ligand binding domain

To understand the molecular basis of its potent estrogenic activity, the X-ray cocrystal structure of **2a** in complex with the ERα ligand binding domain (LBD) was determined. A Y537S mutant of the ERα LBD and a short peptide from the GRIP coactivator, which together favor the agonist conformation of the receptor,^17^ were used to facilitate crystallization. The structure was solved to 1.55 Å using molecular replacement (SI Table 3) and showed a canonical ERα LBD dimer with its C-terminal helix 12 (H12) in the agonist conformation over the ligand binding pocket and with the coactivator GRIP peptide in the cleft that forms the activating function 2 (AF-2) (Figure 3A). Within the ligand binding pocket, the thieno[2,3-*d*]pyrimidine (**2a**) was clearly identified in the electron density map (SI Figure 1). The ligand adopted a binding pose perpendicular to H12 and formed a single hydrogen bond between its phenolic oxygen and E353, but with no additional interaction to R394 or the water molecule that are utilized by E2 (Figure 3B). The 5-phenyl group of **2a** sat adjacent to helix 3 near L384, L387, A350, and L525 and adopted a similar vector to the quaternary methyl group of E2 that is pointed towards H12. The 2-chloro group of **2a** sat in a hydrophobic binding pocket formed by M343, M421, and F425. Both M343 and F425 were displaced away from their conformation in the E2 structure by the presence of the 2-chloro group of **2a**. Although the thieno[2,3-*d*]pyrimidine **2a** induced a canonical agonist conformation where H12 is it folded over the ligand binding pocket, the thiophene ring perturbed the H11-H12 connecting loop and slightly reduced the contact surface of H12 with the ligand binding pocket compared to the E2 structure. The reduced burial of the H12 may explain why **2a** was marginally less efficacious than E2 in the transcriptional activation assay (Figure 2E). Unlike the 2-, 4-, and 5-substituents of the thieno[2,3-*d*]pyrimidine that formed direct contacts with amino acids lining the ERα ligand binding pocket, the polar 1- and 3-pyrimidine nitrogens and the secondary amino group of the 4-substituent did not form any H-bond interactions with the receptor. In comparison to other ERα cocrystal structures, the **2a** complex most closely resembled the phytoestrogen ferutinin (PDB 4MG7), which occupied a similar region of the ligand binding pocket while also forming only a single polar interaction with E353 (Figure 3D).

### Three-step synthesis of 2,4,5-substituted thieno[2,3-d]pyrimidines

To perform a systematic evaluation of the structure-activity of **2a** as a new chemotype of nonsteroidal estrogens, we developed an efficient synthesis of the 2-chlorothieno[2,3-*d*]pyrimidine core that allowed for sequential modification of the 4- and 5-substituents. The first step involved a three-component Gewald reaction to generate trisubstituted thiophene **3a** from acetophenone, malononitrile, and sulfur.^18-19^ Using the one-pot base catalysis conditions reported by Zeng et al,^20-21^ the desired thiophene **3** was isolated, but only in 10% yield (Table 1, entry 1). We performed a systematic optimization of the reaction by varying the base catalyst, solvent, and reaction time. Increase in the reaction temperature to 80 °C and the reaction time to 24 h led to 14% isolated yield of **3a** (entry 2). Increasing the amount of base to 0.25 equivalents of imidazole at the original 60 °C temperature also gave an increase in the yield of **3** to 18 % (entry 3). Switching the solvent to 2-Me-THF (entry 4) resulted in a near doubling of the yield to 32 %. Under these conditions, successive increase in the equivalents of imidazole to 0.5 and 1.0 (entries 5 and 6) gave 36 % and 40 % yields, respectively, of **3**. Increasing the temperature to 80 °C (entry 7) did not give any further increase in yield. Switching the solvent from 2-Me-THF to dioxane (entry 8) resulted in a small decrease in the yield of **3a** to 35%. Some authors have reported use of the bases morpholine and piperidine for catalysis of the Gewald reaction.^18-19^

**Table 1.**
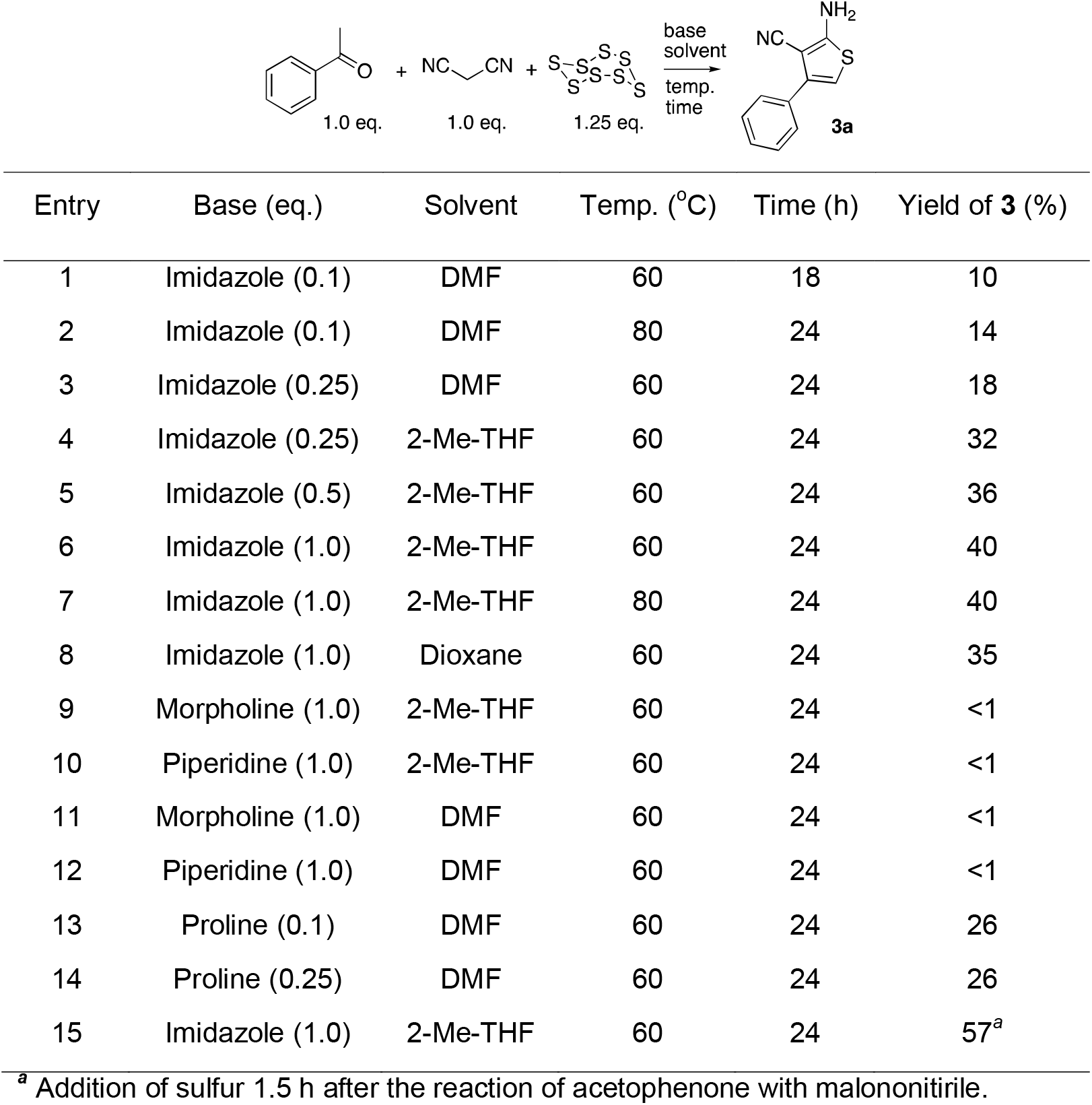
Optimization of the one-pot Gewald reaction

However, in our hands neither of these bases yielded more than traces of **3a** with either 2-Me-THF or DMF as solvent (entries 9-12). Use of proline as the base in DMF^21^ at 0.1 and 0.25 equivalents gave 26% of **3a** (entries 13 and 14) but was still inferior to imidazole. Finally, after consideration of the reaction mechanism for the three-component base catalyzed Gewald reaction (SI Figure 1), which involves the initial formation of Knoevenagel product of malononitrile with acetophenone, we opted to delay addition of the sulfur until formation of the Knoevenagel product was observed by TLC.^18^ Our goal was to minimize the formation of sulfur containing acyclic intermediates that were isolated as various byproducts under the lower yielding reaction conditions (data not shown). Accordingly, when the sulfur addition was delayed for 1.5 h after the mixing of acetophenone and malononitrile with 1.0 equivalent of imidazole in 2-Me-THF at 60 °C, the trisubstituted thiophene **3a** was isolated in 57% yield (entry 15). Having optimized the synthesis of thiophene **3a**, we employed an acetonitrile-assisted diphosgene reaction as reported by Chi^22^ and by Roeker^23^ to form the 2,4-dicholothieno[2,3-*d*]pyrimidine **4a** in high yield (Scheme 1). Finally, **4a** was subjected to S_N_Ar reaction with 4-hydroxybenzylamine to obtain **2a**. Under the optimized conditions, the efficient three-step synthesis thieno[2,3-*d*]pyrimidine **2a** from acetophenone was achieved in 28 % overall yield on a gram scale.

**Scheme 1.**
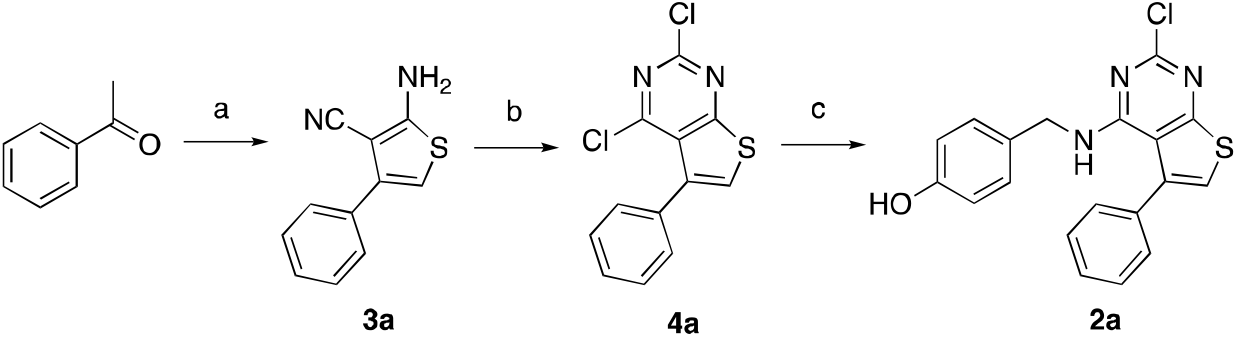
Optimized 3-step synthesis of thienopyrimidine **2a** Reagents and conditions: (a) (i) malononitrile (1.1 eq.), imidazole (1.0 eq.), 2-Me-THF. 60 °C, 1.5 h, then (ii) S_8_ (1.25 eq.), 60 °C, 24 h, 57 %; (b) diphosgene (1.5 eq.), MeCN, sealed tube, 110 °C, 24h, 77 %; (c) 4-hydroxybenylamine, Et_3_N, CHCl_3_, 75 °C, 66%.

### Structure-activity of 2-cholorothieno[2,3-*d*]pyrimidines as ERα agonists

Guided by the cocrystal structure of **2a** in complex with the ERα LBD (Figure 3) we explored the role of the phenolic group at the 4-position and the hydrophobic substituent at the 5-position of the thieno[2,3-*d*]pyrimidine core. The commercially available analogs (SI Table 1) indicated that the 2-chloro substituent was already optimal, so we opted to leave this position constant. Six new analogs (**2b**–**g**) were obtained using the three-step 2-chlorothieno[2,3-*d*]pyrimidine synthesis from the corresponding methyl ketones (Scheme 2). The analogs were tested for their ERα binding affinity using the two-hybrid LiSA and for their agonist functional activity in the ERE-luciferase reporter assay in HepG2 cells. A resynthesized sample of **2a** gave an ERα binding affinity of 11 nM and an EC_50_ = 48 nM in the reporter gene assay (Table 2). The 5-methyl analog **2b** and 5-isopropyl **2c** analog were both >100-fold less active in the receptor binding and reporter gene functional assays. In contrast, the 5-isobutyl analog **2d** was only 3-fold less active than **2a** in the binding assay and 5-fold less active in the reporter gene assay. This result mirrors the structure activity observed in the 2-methylthieno[2,3-*d*]pyrimidines (SI Table 1) where the corresponding 5-isobuyl and 5-phenyl analogs had similar activity on ERα. The 5-tertbutyl-2-chlorothieno[2,3-*d*]pyrimidine **2e** was less active than **2a**. However, the 5-cyclohexyl analog **2f** showed a relatively small decrease in potency, which was notable given the poor activity of the 5-isopropyl analog **2c**. Finally, the analog **2g** with an ortho-chloro substituent on the 5-phenyl group had relatively potent ERα binding and activation, which was consistent with the rotation of the 5-phenyl group out of the plane of the 2-chlorothieno[2,3-*d*]pyrimidine core in the co-crystal structure (Figure 3B). Overall, while the phenyl group remained the optimal 5-substituent, the receptor was able to accommodate branched alkyl substituents and ortho-substituted phenyl groups, which indicated the potential for further optimization of the receptor binding and functional activity.

**Scheme 2.**
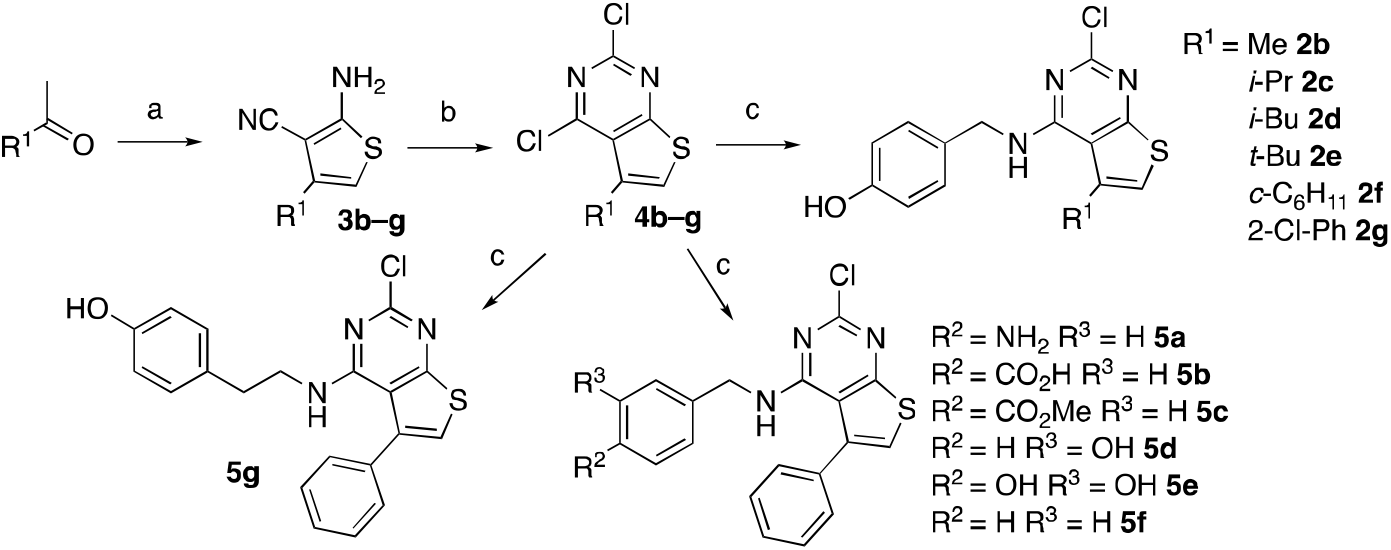
Synthesis of thienopyrimidine analogs **2b**–**g** and **5a**–**g** Reagents and conditions: (a) (i) malononitrile (1.1 eq.), imidazole (1.0 eq.), 2-Me-THF. 60 °C, 1.5 h, then (ii) S_8_ (1.25 eq.), 60 °C, 24 h; (b) diphosgene (1.5 eq.), MeCN, sealed tube, 110 °C, 18-24h; (c) amine, Et_3_N, CHCl_3_*, 75 °C; * for **5b** *t*-BuOH/DIPEA at 130 °C in microwave, for **5e** *t*-BuOH.

**Table 2.** ERα activity of compounds **2a**–**g**

| Compound | R | $\text{ER}\alpha$ Binding<br>$\text{EC}_{50}$ (nM)<sup>a</sup> | $\text{ER}\alpha$ Agonism<br>$\text{EC}_{50}$ (nM)<sup>b</sup> |
| --- | --- | --- | --- |
| **2a** | Ph | 11 | 48 |
| **2b** | Me | 1700 | 6600 |
| **2c** | $i\text{-Pr}$ | 6200 | >10000 |
| **2d** | $i\text{-Bu}$ | 31 | 240 |
| **2e** | $t\text{-Bu}$ | 890 | 2000 |
| **2f** | $c\text{-C}_6\text{H}_{11}$ | 95 | 350 |
| **2g** | 2-Cl-Ph | 43 | 170 |

<sup>a</sup> $\text{ER}\alpha$ LiSA assay; <sup>b</sup> (ERE)<sub>7</sub>-TK-Luc reporter assay in HepG2 cells; i.a. inactive at 10 $\mu\text{M}$ .

A second series of analogs (**5a**–**g**) explored the role of the 4’-phenol in binding to ERα (Table 3). The replacement of the 4’-hydoxyl of **2a** with 4’-amine (**5a**), 4’-carboxylic acid (**5b**), and 4’-methylester group (**3b**) gave analogs that were inactive in the binding and functional assays at 10 µM. Switching from the 4’-phenol of **2a** to the 3’-phenol of **5d** led to a 1000-fold loss in activity. This result was on one hand surprising given the comparison of **2a** with E2 in the X-ray cocrystal structures (Figure 3C), yet was consistent with the observations in the 5-methylthieno[2,3-*d*]pyrimidine analogs (SI Table 1). The detrimental effect of the 3’-phenol was confirmed in the 3’,4’-catechol analog (**5e**) that was inactive in the binding and functional assays. The unsubstituted phenyl analog (**5f**), while retaining sub-µM binding and functional activity, provided quantitative evidence that addition of the 4’-hydroxyl onto the 2-chlorothieno[2,3-*d*]pyrimidine core in **2a** contributed a 20 to 40-fold increase in potency on ERα. Homologation of the linker between the amine and phenol by an extra methylene unit in **5g** resulted in a large decrease in ERα activity.

**Table 3.**
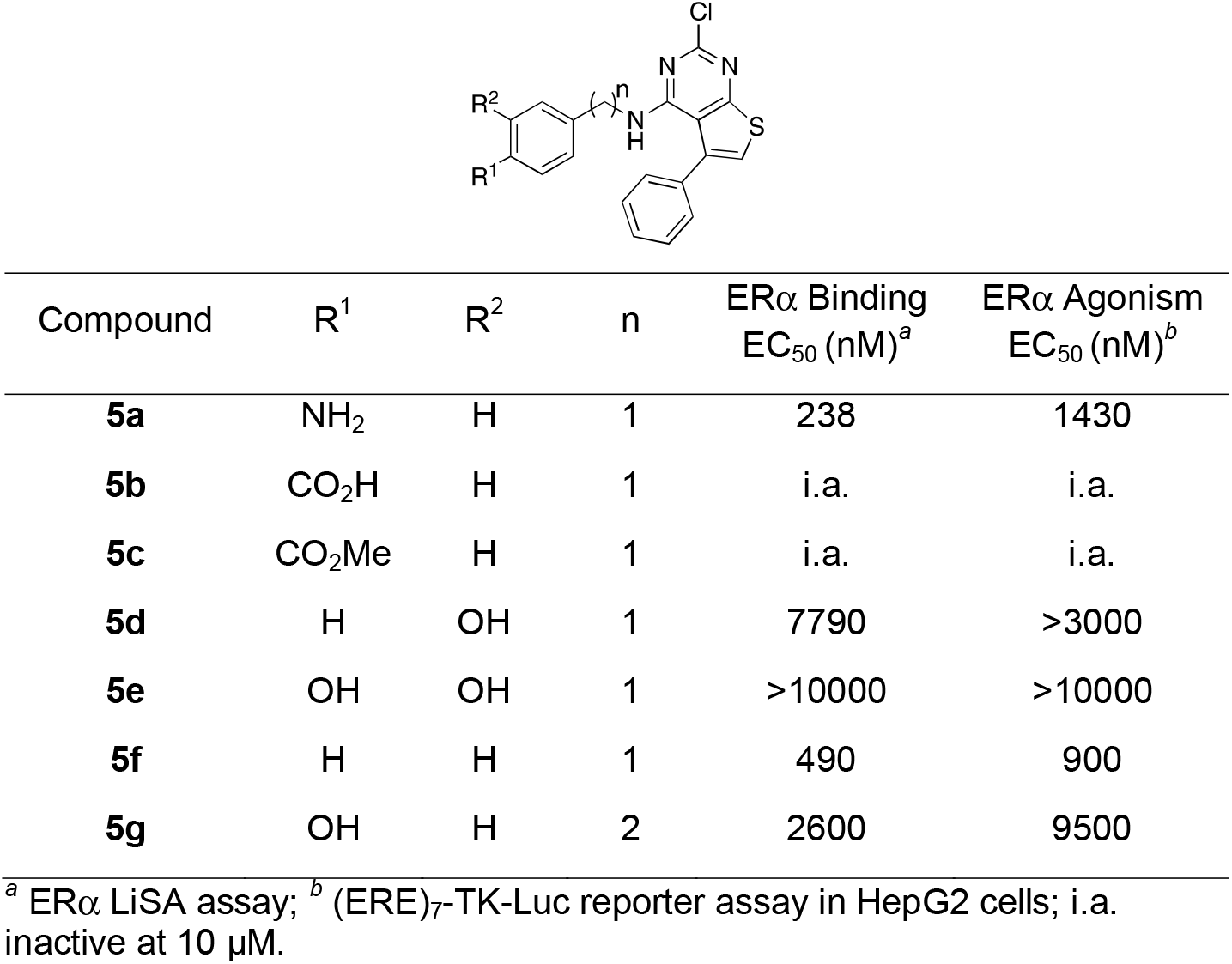
ERα activity of compounds **5a**–**g**

To more closely mimic the binding of E2 to ERα, where the 17β-hydroxyl group forms an H-bond with the H524 residue (Figure 3F),^10^ we sought to add polar functionality at C6 of the thieno[2,3-*d*]pyrimidine core. The C6 substituted analogs were synthesized from 2-methoxyacetophenone using the imidazole catalyzed one-pot Gewald reaction and acetonitrile-assisted diphosgene reactions to give the corresponding 6-methoxy-2,4-dichlorothieno[2,3-*d*]pyrimidine (Scheme 3). S_N_Ar reaction with 4-hydroxybenzylamine yielded the 6-methoxy analog **6a**. Demethylation with BBr_3_ produced a mixture of 6-hydroxythieno[2,3-*d*]pyrimidine **6b** and its keto tautomer **6b’**. The ratio of **6b’**:**6b** favored the keto form by a 9:1 ratio in d_6_-DMSO as determined by the ^1^H-NMR assignment (SI Figure 2), and it is likely that they readily interconvert in protic solvents. When tested in the ERα LISA (Table 4), the 6-methoxy analog (**6a**) was 10-fold less potent than the unsubstituted thieno[2,3-*d*]pyrimidine (**2a**). The 6-hydroxy/keto analog (**6b**/**6b’**) showed improved potency compared to **6a** with EC_50_ = 40 nM, although it was 3-fold less potent than **2a** and also showed slightly lower binding affinity (Figure 2d). Notably, **6b/6b’** had equivalent potency to **2a** in the ERE reporter assay. X-ray crystallography was used to determine the preferred tautomer of **6b**/**6b’** for binding to ERα (Figure 3E and SI Table 3). The electron density map (SI Figure 1) clearly showed the enol form **6b** in the binding pocket, with an sp2 center at C-5 on the thieno[2,3-*d*]pyrimidine and the 6-hydroxyl group forming an H-bond with H524. The crystal structure showed that thieno[2,3-*d*]pyrimidine (**6b**) was able to engage both of the polar residues (E353 and H524) that are utilized by the natural hormone E2 for molecular recognition of ERα. The small decrease in potency of **6b**/**6b’** compared to unsubstituted **2a** in the ERα LiSA and binding assay may be due to the ratio of enol:keto tautomers in the assay media. It is notable that estrone, the 17-keto analog of E2 (Figure 1), is 10-100-fold less potent as a hormone,^24^ consistent with a preference for an H-bond donor over an acceptor in the interaction with H524.

**Table 4.**
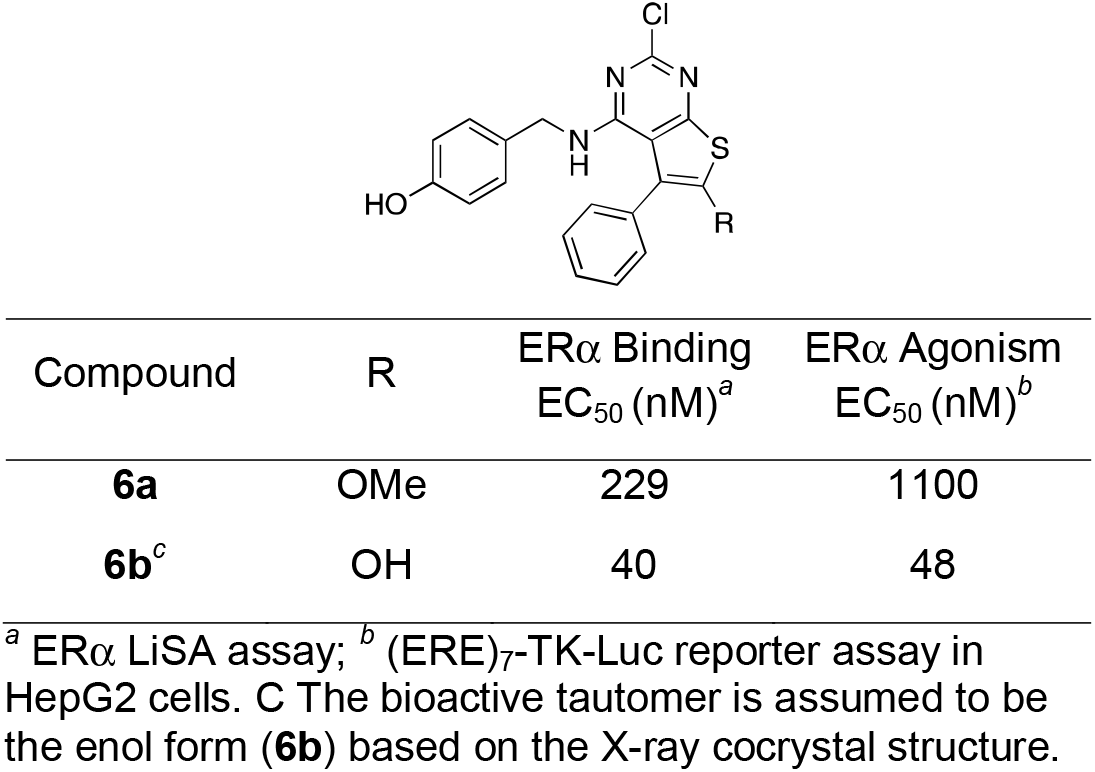
ERα activity of compounds **6a** and **6b**

| Compound | R | ERα Binding<br>EC<sub>50</sub> (nM)<sup>a</sup> | ERα Agonism<br>EC<sub>50</sub> (nM)<sup>b</sup> |
| --- | --- | --- | --- |
| **6a** | OMe | 229 | 1100 |
| **6b<sup>c</sup>** | OH | 40 | 48 |

<sup>a</sup> ERα LiSA assay; <sup>b</sup> (ERE)<sub>7</sub>-TK-Luc reporter assay in HepG2 cells. <sup>c</sup> The bioactive tautomer is assumed to be the enol form (**6b**) based on the X-ray cocrystal structure.

## CONCLUSIONS

2-chloro-4-((4-hydroxybenzyl)amino)-5-phenylthieno[2,3-d]pyrimidin-6-ol (**6b**) exemplifies a new chemotype of potent nonsteroidal estrogens. While there are no prior reports of thieno[2,3-*d*]pyrimidines with activity as steroid receptor ligands, these and related heterocycles have been identified as ATP-competitive kinase inhibitors.^25-26^ However, when screened, at a concentration of 20 µM, across a panel of 100 human protein kinases **2a** showed little or no effect on protein thermal stability^27^ (SI Table 4), indicating that it is unlikely to have significant kinase inhibitory activity. Importantly, thieno[2,3-*d*]pyrimidines are synthetically accessible through a modular three-step synthesis (Schemes 1-3) that supports chemical modification at each position on the heterocyclic core. Furthermore, X-ray crystallography revealed the detailed interactions of **2a** and **6b** within the ERα ligand binding pocket and identified the molecular interactions contributing to their potency and efficacy as estrogens. These molecular insights combined with the chemical tractability of the thieno[2,3-*d*]pyrimidine chemotype will support the design and synthesis of a new nonsteroidal estrogen agonists, partial agonists, and antagonists as potential therapeutic hormones and antihormones.

## EXPERIMENTAL SECTION

### General Procedures

All the reactions were performed in oven-dried glassware. Thin-layer chromatography (TLC) was performed using aluminum backed silica coated plates. TLC plates were visualized under Ultraviolet light. The ^1^H and ^13^C NMRs were recorded on Agilent 400MR INOVA NMR, [^1^H (400 MHz), ^13^C (101 MHz)] and Bruker Avance III HD 850 MHz four channel spectrometer equipped with a TCI H-C/N-D 5 mm CryoProbe [^1^H (850 MHz), ^13^C (214 MHz)] with complete proton decoupling for ^13^C. Chemical shifts were analyzed on MestReNova software. Chemical shifts are reported in parts per million with the solvent resonance as the internal standard (CDCl_3_, ^1^H: δ7.26 ppm, ^13^C: δ77.16 ppm; DMSO-*d*^6, 1^H: δ2.50 ppm, ^13^C: δ39.52 ppm). Coupling constants are reported in Hertz (Hz). Abbreviations are used as follows: s = singlet, d = doublet, t = triplet, q = quartet, hept = heptet, m = multiplet, dd=doublet of doublet. Samples were analyzed with a ThermoFisher Q Exactive HF-X (ThermoFisher, Bremen, Germany) mass spectrometer (electrospray ionization) coupled with a Waters Acquity H-class liquid chromatograph system.

### Procedure for the synthesis of thiophene intermediates 3a–h

To a reaction vial containing corresponding methyl ketone (3 mmol, 1 equiv) in dry methyl-THF (5 mL) was added malononitrile (1.1 equiv) followed by imidazole (1.0 equiv) at room temperature, the mixture was allowed to stir under nitrogen at 60°C for 1.5 to 2h. Sulfur (1.25 equiv) was then added to the reaction mixture in one portion and the contents were stirred at 60°C for 22.5h.The reaction contents were then cooled to room temperature followed by transfer into a separatory funnel containing 50 mL water. Ethyl acetate (30 mL) was then added to the separatory funnel and the extraction was performed. The aqueous layer was washed with 10 mL ethyl acetate. The combined organic layers were washed with water followed by brine, dried over anhydrous sodium sulfate and evaporated to yield crude residue that was purified through flash chromatography by gradient elution (Ethyl acetate : Hexane).

*2-amino-4-phenylthiophene-3-carbonitrile (***3a**): Light yellow solid; Yield 57%; ^1^H NMR (850 MHz, DMSO-*d*_6_) δ 7.55 (d, J = 7.3 Hz, 2H), 7.44 (t, J = 7.7 Hz, 2H), 7.37 (t, J = 7.4 Hz, 1H), 7.23 (s, 2H), 6.54 (s, 1H); ^13^C NMR (214 MHz, DMSO-*d*_6_) δ 169.6, 141.6, 137.6, 131.8, 130.9, 130.0, 119.7, 108.19, 86.4. ; HRMS (ES+) m/z calc. for [C_11_H_8_N_2_S + H^+^]: 201.0481; found: 201.0479.

*2-amino-4-methylthiophene-3-carbonitrile (***3b**): Light yellow solid: Yield 37%; ^1^H NMR (400 MHz, DMSO-*d*_6_) δ 7.03 (s, 2H), 6.01 (s, 1H), 2.02 (s, 3H); ^13^C NMR (101 MHz, DMSO-*d*_6_) δ 165.4, 134.7, 116.7, 103.7, 86.0, 15.5; HRMS (ES+) m/z calc. for [C_6_H_6_N_2_S + H^+^]: 139.0324; found: 139.0324.

*2-amino-4-isopropylthiophene-3-carbonitrile (***3c**): Brown solid; Yield 39%; ^1^H NMR (400 MHz, DMSO-*d*_6_) δ 7.00 (s, 2H), 5.99 (s, 1H), 2.71 (hept, J = 7.2 Hz, 1H), 1.15 (d, J = 6.8 Hz, 6H); ^13^C NMR (101 MHz, DMSO-*d*_6_) δ 165.9, 146.2, 116.8, 101.4, 84.6, 29.3, 22.6; HRMS (ES+) m/z calc. for [C_8_H_10_N_2_S + H^+^]: 167.0637; found: 167.0636.

*2-amino-4-isobutylthiophene-3-carbonitrile (***3d**): Light brown glue; Yield 41% ; ^1^H NMR (850 MHz, DMSO-*d*_6_) δ 7.03 (s, 2H), 6.89 (s, 1H), 6.03 (s, 1H), 3.09 (hept, *J* = 6.8 Hz, 1H), 2.27 (d, *J* = 7.2 Hz, 2H), 1.97 (s, 2H), 1.86 (dh, *J* = 13.6, 6.8 Hz, 1H), 1.11 (d, *J* = 6.8 Hz, 4H), 0.88 (d, *J* = 6.6 Hz, 6H); ^13^C NMR (214 MHz, DMSO-*d*_6_) δ 168.1, 164.8, 141.3, 130.3, 129.7, 119.7, 119.5, 106.8, 88.5, 88.4, 41.7, 31.0, 30.0, 27.3, 25.3, 15.5; HRMS (ES+) m/z calc. for [C_9_H_12_N_2_S + H^+^]: 181.0794; found: 181.0792.

*2-amino-4-(tert-butyl)thiophene-3-carbonitrile (****3e***): Cream color solid; Yield 19%; ^1^H NMR (400 MHz, DMSO-*d*_6_) δ 6.95 (s, 2H), 5.97 (s, 1H), 1.24 (s, 9H); ^13^C NMR (101 MHz, DMSO-*d*_6_) δ 167.1, 148.4, 117.9, 101.5, 83.6, 34.1, 29.7; HRMS (ES+) m/z calc. for [C_9_H_12_N_2_S + H^+^]: 181.0794; found: 181.0792.

*2-amino-4-cyclohexylthiophene-3-carbonitrile (***3f**): Light yellow solid; Yield 47%; ^1^H NMR (400 MHz, DMSO-*d*_6_) δ 6.99 (s, 2H), 5.98 (s, 1H), 2.41 – 2.27 (m, 1H), 1.88 – 1.60 (m, 5H), 1.28 (t, *J* = 10.3 Hz, 4H), 1.20 – 1.07 (m, 1H); ^13^C NMR (101 MHz, DMSO-*d*_6_) δ 165.7, 145.4, 116.8, 101.6, 84.8, 39.2, 32.9, 26.5, 26.1; HRMS (ES+) m/z calc. for [C_11_H_14_N_2_S + H^+^]: 207.0950; found: 207.0948.

*2-amino-4-(2-chlorophenyl)thiophene-3-carbonitrile (***3g**): Yellow solid; Yield 54%; ^1^H NMR (850 MHz, DMSO-*d*_6_) δ 7.55 (dd, *J* = 7.8, 1.3 Hz, 1H), 7.44 – 7.37 (m, 3H), 7.24 (s, 2H), 6.44 (s, 1H); ^13^C NMR (214 MHz, DMSO-*d*_6_) δ 168.2, 139.2, 136.9, 135.3, 134.6, 133.0, 132.8, 130.4, 119.0, 110.4, 88.5; HRMS (ES+) m/z calc. for [C_11_H_7_ClN_2_S + H^+^]: 235.0091; found: 235.0090.

*2-amino-5-methoxy-4-phenylthiophene-3-carbonitrile (***3h**): Brown to red solid; Yield 75%; ^1^H NMR (850 MHz, DMSO-*d*_6_) δ 7.45 – 7.41 (m, 4H), 7.33 (tt, *J* = 6.87, 2.05 Hz, 1H), 7.00 (s, 2H), 3.70 (s, 3H); ^13^C NMR (214 MHz, DMSO-*d*_6_) δ 159.8, 146.5, 135.1, 131.5, 131.5, 130.4, 122.2, 119.6, 83.0, 66.2; HRMS (ES+) m/z calc. for [C_12_H_10_N_2_OS + H^+^]: 231.0587; found: 231.0583.

### Procedure for the synthesis of thienopyrimidine intermediates 4a–g

To a microwave reaction vial containing corresponding thiophene (**3a - 3g**, 1 mmol, 1 equiv) in dry acetonitrile (4 mL) was added diphosgene (1.5 equiv) at room temperature under nitrogen atmosphere, a white precipitate occurred (within 10 min of stirring at room temperature). The mixture was then allowed to stir under nitrogen at 110°C until the disappearance of starting material (per TLC, typical reaction times were between 18h – 24h). The reaction contents were then cooled to room temperature followed by transfer into a separatory funnel containing 30 mL water. Ethyl acetate (20 mL) was then added to the separatory funnel and the extraction was performed. The aqueous layer was washed with 10 mL ethyl acetate. The combined organic layers were washed with brine, dried over anhydrous sodium sulfate and evaporated to yield crude residue that was purified through flash chromatography by gradient elution (Ethyl acetate : Hexane).

*2,4-dichloro-5-phenylthieno[2,3-d]pyrimidine (***4a**): White solid; Yield 77%; ^1^H NMR (850 MHz, DMSO-*d*_6_) δ 8.02 (s, 1H), 7.49 – 7.47 (m, 5H); ^13^C NMR (214 MHz, DMSO-*d*_6_) δ 174.2, 158.2, 155.7, 137.7, 137.4, 133.1, 131.7, 131.5, 131.0, 128.9; HRMS (ES+) m/z calc. for [C_11_H_7_ClN_2_S + H^+^]: 280.9702; found: 280.9699.

*2,4-dichloro-5-methylthieno[2,3-d]pyrimidine (***4b**): White solid; Yield 49%; ^1^H NMR (850 MHz, DMSO-*d*_6_) δ 7.78 (q, *J* = 1.4 Hz, 1H), 2.61 (d, *J* = 1.4 Hz, 3H); ^13^C NMR (214 MHz, DMSO-*d*_6_) δ 174.7, 158.2, 155.6, 132.8, 130.0, 129.1, 20.1; HRMS (ES+) m/z calc. for [C_7_H_4_Cl_2_N_2_S + H^+^]: 218.9545; found: 218.9540.

*2,4-dichloro-5-isopropylthieno[2,3-d]pyrimidine (***4c**): White solid; Yield 62%; ^1^H NMR (400 MHz, DMSO-*d*_6_) δ 7.81 (s, 1H), 3.68 (hept, *J* = 6.6 Hz, 1H), 1.30 (d, *J* = 6.8 Hz, 6H); ^13^C NMR (101 MHz, DMSO-*d*_6_) δ 172.4, 154.9, 152.8, 141.4, 126.4, 123.9, 28.6, 23.7; HRMS (ES+) m/z calc. for [C_9_H_8_Cl_2_N_2_S + H^+^]: 246.9858; found: 246.9854.

*2,4-dichloro-5-isobutylthieno[2,3-d]pyrimidine (***4d**): Colorless glue; Yield 45%; ^1^H NMR (400 MHz, Chloroform-*d*) δ 7.20 (s, 1H), 3.47 (hept, *J* = 6.7 Hz, 1H), 2.89 (d, *J* = 7.0 Hz, 2H), 2.57 (s, 2H), 1.99 (hept, *J* = 6.7 Hz, 1H), 1.35 (d, *J* = 6.8 Hz, 5H), 0.98 (d, *J* = 6.6 Hz, 6H); ^13^C NMR (101 MHz, Chloroform-*d*) δ 172.3, 169.8, 155.5, 154.3, 153.7, 152.7, 150.3, 134.3, 128.2, 126.4, 124.4, 123.0, 39.7, 29.1, 28.4, 23.9, 22.1, 14.2; HRMS (ES+) m/z calc. for [C_10_H_10_Cl_2_N_2_S + H^+^]: 261.0015; found: 261.0011.

*5-(tert-butyl)-2,4-dichlorothieno[2,3-d]pyrimidine (***4e**): White solid; Yield 33%; ^1^H NMR (850 MHz, DMSO-*d*_6_) δ 7.88 (s, 1H), 1.55 (s, 9H); ^13^C NMR (214 MHz, DMSO-*d*_6_) δ 177.1, 157.1, 155.1, 146.2, 129.8, 127.7, 37.6, 34.3; HRMS (ES+) m/z calc. for [C_10_H_10_Cl_2_N_2_S + H^+^]: 261.0015; found: 261.0011.

*2,4-dichloro-5-cyclohexylthieno[2,3-d]pyrimidine (***4f**): White solid; Yield 87%; ^1^H NMR (850 MHz, DMSO-*d*_6_) δ 7.79 (d, *J* = 0.9 Hz, 1H), 3.30 (td, *J* = 11.1, 2.6 Hz, 1H), 2.02 (d, *J* = 10.9 Hz, 2H), 1.83 (d, *J* = 12.6 Hz, 2H), 1.75 (d, *J* = 13.2 Hz, 1H), 1.48 – 1.38 (m, 4H), 1.26 (dddd, *J* = 16.5, 12.8, 8.5, 3.8 Hz, 1H); ^13^C NMR (214 MHz, DMSO-*d*_6_) δ 175.0, 157.7, 155.4, 143.1, 129.0, 126.8, 41.2, 36.8, 29.3, 28.7; HRMS (ES+) m/z calc. for [C_12_H_12_Cl_2_N_2_S + H^+^]: 287.0171; found: 287.0170.

*2,4-dichloro-5-(2-chlorophenyl)thieno[2,3-d]pyrimidine (***4g**): White solid; Yield 84%; ^1^H NMR (850 MHz, DMSO-*d*_6_) δ 8.12 (s, 1H), 7.62 (d, *J* = 8.1 Hz, 1H), 7.54 (td, *J* = 7.7, 1.8 Hz, 1H), 7.51 (dd, *J* = 7.5, 1.8 Hz, 1H), 7.48 (td, *J* = 7.4, 1.2 Hz, 1H); ^13^C NMR (214 MHz, DMSO-*d*_6_) δ 173.8, 158.1, 156.1, 136.8, 136.5, 135.3, 134.2, 133.9, 132.8, 132.2, 130.3, 129.1; HRMS (ES+) m/z calc. for [C_12_H_5_Cl_3_N_2_S + H^+^]: 314.9312; found: 314.9307.

*2,4-dichloro-6-methoxy-5-phenylthieno[2,3-d]pyrimidine (***4h**): White solid; Yield 70%; ^1^H NMR (850 MHz, DMSO-*d*_6_) δ 7.47 – 7.43 (m, 2H), 7.43 – 7.40 (m, 1H), 7.40 – 7.38 (m, 2H), 4.06 (s, 3H); ^13^C NMR (214 MHz, DMSO-*d*_6_) δ 165.7, 164.6, 154.7, 153.3, 134.4, 134.2, 131.2, 131.0, 129.8, 114.8, 65.9; HRMS (ES+) m/z calc. for [C_13_H_8_Cl_2_N_2_OS + H^+^]: 310.9807; found: 310.9802.

### Procedure for the synthesis of compounds 2a–g and 6a

To a reaction vial containing 4-hydroxybenzylamine (∼30 mg, 0.24 mmol, 1.2 equiv) was added triethylamine (5 equiv) dropwise in one portion and the contents were stirred at 75°C until a clear solution is obtained. To the transparent solution corresponding thienopyrimidine (**4a**–**g**, 0.2 mmol, 1 equiv) was added in one portion and stirred at 75°C until the disappearance of thienopyrimidne (per TLC, typical reaction time was ∼24h). The reaction contents were then cooled to room temperature followed by transfer into a separatory funnel containing 30 mL water. DCM (25 mL) was then added to the separatory funnel and the extraction was performed. The organic layer was washed with brine, dried over anhydrous sodium sulfate and evaporated to yield crude residue that was purified through flash chromatography by gradient elution (Ethyl acetate : Hexane).

*4-(((2-chloro-5-phenylthieno[2,3-d]pyrimidin-4-yl)amino)methyl)phenol (***2a**): White Solid; Yield 66%; ^1^H NMR (400 MHz, DMSO-*d*_6_) δ 9.33 (s, 1H), 7.51 (s, 1H), 7.48 – 7.37 (m, 5H), 6.98 (d, *J* = 8.5 Hz, 2H), 6.65 (d, *J* = 8.6 Hz, 2H), 5.81 (t, *J* = 5.2 Hz, 1H), 4.39 (d, *J* = 5.2 Hz, 2H); ^13^C NMR (101 MHz, DMSO-*d*_6_) δ 167.7, 157.9, 157.0, 155.4, 135.2, 134.7, 129.5, 129.4, 129.2, 129.0, 127.9, 122.1, 115.6, 112.7, 44.4; HRMS (ES+) m/z calc. for [C_19_H_14_ClN_3_OS + H^+^]: 368.0619; found: 368.0614.

*4-(((2-chloro-5-methylthieno[2,3-d]pyrimidin-4-yl)amino)methyl)phenol (***2b**): White solid; Yield 54%; ^1^H NMR (850 MHz, DMSO-*d*_6_) δ 9.28 (s, 1H), 7.59 (t, *J* = 5.90 Hz, 1H), 7.20 (s, 1H), 7.19 (d, *J* = 5.87 Hz, 2H), 6.71 (d, *J* = 8.54 Hz, 2H), 4.59 (d, *J* = 5.77 Hz, 2H), 2.58 (s, 3H); ^13^C NMR (214 MHz, DMSO-*d*_6_) δ 170.9, 161.4, 159.4, 157.8, 132.6, 132.2, 131.8, 121.5, 118.1, 118.1, 117.5, 46.6, 20.3; HRMS (ES+) m/z calc. for [C_14_H_12_ClN_3_OS + H^+^]: 306.0462; found: 306.0458.

*4-(((2-chloro-5-isopropylthieno[2,3-d]pyrimidin-4-yl)amino)methyl)phenol (***2c**): White solid; Yield 71%; ^1^H NMR (850 MHz, DMSO-*d*_6_) δ 9.28 (s, 1H), 7.57 (t, *J* = 5.9 Hz, 1H), 7.24 (s, 1H), 7.20 (d, *J* = 8.5 Hz, 2H), 6.70 (d, *J* = 8.5 Hz, 2H), 4.61 (d, *J* = 5.8 Hz, 2H), 3.50 (hept, *J* = 6.5 Hz, 1H), 1.25 (d, *J* = 6.6 Hz, 6H); ^13^C NMR (214 MHz, DMSO-*d*_6_) δ 171.4, 160.8, 159.4, 157.6, 144.4, 132.3, 132.0, 118.6, 118.1, 116.4, 46.8, 31.4, 26.3; HRMS (ES+) m/z calc. for [C_16_H_16_ClN_3_OS + H^+^]: 334.0775; found: 334.0769.

*4-(((2-chloro-5-isobutylthieno[2,3-d]pyrimidin-4-yl)amino)methyl)phenol (***2d**): White solid; Yield 38%; ^1^H NMR (400 MHz, DMSO-*d*_6_) δ 9.28 (s, 1H), 7.29 (t, *J* = 5.7 Hz, 1H), 7.16 (d, *J* = 8.6 Hz, 3H), 6.69 (d, *J* = 8.5 Hz, 2H), 4.57 (d, *J* = 5.6 Hz, 2H), 2.80 (d, *J* = 6.9 Hz, 2H), 1.80 (dp, *J* = 13.4, 6.8 Hz, 1H), 0.84 (d, *J* = 6.6 Hz, 6H); ^13^C NMR (101 MHz, DMSO-*d*_6_) δ 168.5, 158.4, 156.8, 154.9, 133.8, 129.3, 129.2, 119.5, 115.5, 114.2, 44.1, 28.6, 22.4; HRMS (ES+) m/z calc. for [C_17_H_18_ClN_3_OS + H^+^]: 348.0932; found: 348.0927.

*4-(((5-(tert-butyl)-2-chlorothieno[2,3-d]pyrimidin-4-yl)amino)methyl)phenol (***2e**): White solid; Yield 55%; ^1^H NMR (850 MHz, DMSO-*d*_6_) δ 9.35 (s, 1H), 7.33 (s, 1H), 7.24 (d, *J* = 8.6 Hz, 2H), 6.73 (d, *J* = 8.6 Hz, 2H), 6.62 (t, *J* = 5.5 Hz, 1H), 4.69 (d, *J* = 5.4 Hz, 2H), 1.42 (s, 9H); ^13^C NMR (214 MHz, DMSO-*d*_6_) δ 173.0, 160.0, 159.7, 157.3, 147.0, 132.3, 131.6, 120.0, 118.3, 116.7, 47.7, 36.7, 34.3; HRMS (ES+) m/z calc. for [C_17_H_18_ClN_3_OS + H^+^]: 348.0932; found: 348.0928.

*4-(((2-chloro-5-cyclohexylthieno[2,3-d]pyrimidin-4-yl)amino)methyl)phenol (***2f**): White solid; Yield 52%; ^1^H NMR (400 MHz, DMSO-*d*_6_) δ 9.29 (s, 1H), 7.27 (t, *J* = 5.7 Hz, 1H), 7.17 (d, *J* = 8.9 Hz, 3H), 6.69 (d, *J* = 8.5 Hz, 2H), 4.59 (d, *J* = 5.5 Hz, 2H), 3.13 (t, *J* = 11.2 Hz, 1H), 1.92 (d, *J* = 12.6 Hz, 2H), 1.69 (d, *J* = 10.8 Hz, 3H), 1.56 – 1.37 (m, 2H), 1.33 – 1.09 (m, 3H); ^13^C NMR (101 MHz, DMSO-*d*_6_) δ 168.6, 158.2, 156.8, 154.8, 140.8, 129.3, 129.2, 116.4, 115.5, 113.8, 44.2, 37.9, 33.9, 26.1, 25.8; HRMS (ES+) m/z calc. for [C_19_H_20_ClN_3_OS + H^+^]: 374.1088; found: 374.1083.

*4-(((2-chloro-5-(2-chlorophenyl)thieno[2,3-d]pyrimidin-4-yl)amino)methyl)phenol (***2g**): White solid; Yield 77%; ^1^H NMR (850 MHz, DMSO-*d*_6_) δ 9.34 (s, 1H), 7.61 – 7.57 (m, 2H), 7.53 (dd, *J* = 7.4, 1.7 Hz, 1H), 7.48 (td, *J* = 7.8, 1.8 Hz, 1H), 7.44 (td, *J* = 7.5, 1.3 Hz, 1H), 6.94 (d, *J* = 8.6 Hz, 2H), 6.66 (d, *J* = 8.4 Hz, 2H), 5.51 (t, *J* = 5.4 Hz, 1H), 4.44 (dd, *J* = 14.6, 5.6 Hz, 1H), 4.34 (dd, *J* = 14.6, 5.1 Hz, 1H); ^13^C NMR (214 MHz, DMSO-*d*_6_) δ 169.9, 160.6, 159.7, 158.2, 136.4, 136.0, 135.2, 134.1, 133.7, 133.0, 131.6, 131.0, 130.6, 125.9, 118.3, 116.2, 47.1; HRMS (ES+) m/z calc. for [C_19_H_13_Cl_2_N_3_OS + H^+^]: 402.0229; found: 402.0225.

*4-(((2-chloro-6-methoxy-5-phenylthieno[2,3-d]pyrimidin-4-yl)amino)methyl)phenol (***6a**): Light yellow solid; Yield 63%; ^1^H NMR (400 MHz, DMSO-*d*_6_) δ 9.32 (s, 1H), 7.69 – 7.18 (m, 9H), 6.88 (d, *J* = 8.46 Hz, 3H), 6.63 (d, *J* = 8.51 Hz, 2H), 5.29 (t, *J* = 5.31 Hz, 1H), 4.31 (d, *J* = 4.96 Hz, 3H), 3.89 (s, 3H); ^13^C NMR (101 MHz, DMSO-*d*_6_) δ 157.6, 157.0, 157.0, 156.8, 153.4, 132.0, 130.7, 129.3, 129.0, 128.8, 127.9, 115.6, 112.7, 112.6, 62.9, 44.4; HRMS (ES+) m/z calc. for [C_20_H_16_ClN_3_O_2_S + H^+^]: 398.0725; found: 398.0720.

### Procedure for the synthesis of compounds 5a–5g

For compounds **5a, 5c**, and **5d**–**5g**: To a reaction vial containing substituted benzylamine (1.2 equiv, 0.24 mmol) was added triethylamine (5 equiv) dropwise in one portion and the contents were stirred at 75°C until a clear solution is obtained (0.5 mL methanol was added in case of no clear solution in ∼5 min). To the transparent solution, thienopyrimidine **4a** (0.2 mmol, 1 equiv) was added in one portion and stirred at 75°C until the disappearance of thienopyrimidne (per TLC, typical reaction times were ∼24h). The reaction contents were then cooled to room temperature followed by transfer into a separatory funnel containing 30 mL water. DCM (25 mL) was then added to the separatory funnel and the extraction was performed. The organic layer was washed with brine, dried over anhydrous sodium sulfate and evaporated to yield crude residue that was purified through flash chromatography by gradient elution (Ethyl acetate : Hexane). For compound **5g**, *t*- BuOH was used as solvent with rest of the procedure same as above. For compound **5b**: To a microwave reaction vial containing 4-(aminomethyl)benzoic acid (36 mg, 1.2 equiv, 0.24 mmol) was added DIPEA (2 equiv, 0.4 mmol) followed by thienopyrimidine **4a** (0.2 mmol, 1 equiv) and the contents were stirred at 130 °C for ∼1h under microwave conditions. The reaction mixture was evaporated to yield crude residue that was purified through flash chromatography by gradient elution (DCM : MeOH).

*N-(4-aminobenzyl)-2-chloro-5-phenylthieno[2,3-d]pyrimidin-4-amine (***5a**): Light yellow solid; Yield 87%; ^1^H NMR (850 MHz, DMSO-*d*_6_) δ 7.53 (s, 1H), 7.50 – 7.39 (m, 5H), 6.83 (d, *J* = 8.33 Hz, 2H), 6.47 (d, *J* = 8.38 Hz, 2H), 5.73 (t, *J* = 5.25 Hz, 1H), 5.03 (s, 2H), 4.34 (d, *J* = 5.13 Hz, 2H); ^13^C NMR (214 MHz, DMSO-*d*_6_) δ 170.3, 160.5, 158.1, 151.1, 137.9, 137.4, 132.2, 132.0, 131.7, 131.5, 127.0, 124.7, 116.9, 116.8, 115.4, 47.4; HRMS (ES+) m/z calc. for [C_19_H_15_ClN_4_S + H^+^]: 367.0779; found: 367.0774.

*4-(((2-chloro-5-phenylthieno[2,3-d]pyrimidin-4-yl)amino)methyl)benzoic acid (***5b**): White solid; Yield 43%; ^1^H NMR (400 MHz, DMSO-*d*_6_) δ 7.86 (d, *J* = 8.07 Hz, 2H), 7.57 – 7.36 (m, 6H), 7.31 (d, *J* = 8.18 Hz, 2H), 6.23 (t, *J* = 5.23 Hz, 1H), 4.62 (d, *J* = 5.70 Hz, 2H).; ^13^C NMR (101 MHz, DMSO-*d*_6_) δ 168.0, 167.6, 158.2, 155.2, 143.5, 135.2, 134.9, 130.2, 129.9, 129.5, 129.4, 129.1, 127.6, 122.4, 112.8, 44.4; HRMS (ES+) m/z calc. for [C_20_H_14_ClN_3_O_2_S + H^+^]: 396.0568; found: 396.0565.

*Methyl 4-(((2-chloro-5-phenylthieno[2,3-d]pyrimidin-4-yl)amino)methyl)benzoate (***5c**): White solid; Yield 55%; ^1^H NMR (400 MHz, DMSO-*d*_6_) δ 7.88 (d, *J* = 8.31 Hz, 2H), 7.56 – 7.38 (m, 6H), 7.35 (d, *J* = 8.19 Hz, 2H), 6.27 (t, *J* = 5.44 Hz, 1H), 4.63 (d, *J* = 5.79 Hz, 2H), 3.82 (s, 3H); ^13^C NMR (101 MHz, DMSO-*d*_6_) δ 168.0, 166.5, 158.2, 155.2, 144.2, 135.2, 134.9, 129.7, 129.5, 129.4, 129.1, 128.8, 127.9, 122.4, 112.8, 52.6, 44.4; HRMS (ES+) m/z calc. for [C_21_H_16_ClN_3_O_2_S + H^+^]: 410.0725; found: 410.0721.

*3-(((2-chloro-5-phenylthieno[2,3-d]pyrimidin-4-yl)amino)methyl)phenol (****5d***): White solid; Yield 58%; ^1^H NMR (400 MHz, DMSO-*d*_6_) δ 9.34 (s, 1H), 7.53 (s, 1H), 7.51 – 7.38 (m, 5H), 7.07 (t, *J* = 7.67 Hz, 1H), 6.59 (t, *J* = 11.81 Hz, 3H), 5.98 (t, *J* = 5.39 Hz, 1H), 4.46 (d, *J* = 5.53 Hz, 2H); ^13^C NMR (101 MHz, DMSO-*d*_6_) δ 167.9, 158.1, 157.9, 155.4, 139.5, 135.2, 134.8, 129.9, 129.5, 129.3, 129.1, 122.2, 118.1, 114.6, 114.5, 112.7, 44.6; HRMS (ES+) m/z calc. for [C_19_H_14_ClN_3_OS + H^+^]: 368.0619; found: 368.0615.

*4-(((2-chloro-5-phenylthieno[2,3-d]pyrimidin-4-yl)amino)methyl)benzene-1,2-diol (****5e***): Light brown solid; Yield 59%; ^1^H NMR (400 MHz, DMSO-*d*_6_) δ 8.84 (s, 1H), 8.80 (s, 1H), 7.52 (s, 1H), 7.48 – 7.37 (m, 5H), 6.61 (d, *J* = 8.02 Hz, 1H), 6.58 (d, *J* = 2.05 Hz, 1H), 6.40 (dd, *J* = 7.95, 1.99 Hz, 1H), 5.80 (t, *J* = 5.29 Hz, 1H), 4.34 (d, *J* = 5.20 Hz, 2H); ^13^C NMR (101 MHz, DMSO-*d*_6_) δ 167.9, 158.1, 157.9, 155.4, 139.5, 135.2, 134.8, 129.9, 129.5, 129.3, 129.1, 122.2, 118.1, 114.6, 114.5, 112.7, 44.6; HRMS (ES+) m/z calc. for [C_19_H_14_ClN_3_O_2_S + H^+^]: 384.0568; found: 384.0562.

*N-benzyl-2-chloro-5-phenylthieno[2,3-d]pyrimidin-4-amine (****5f***): White solid; Yield 54%; ^1^H NMR (400 MHz, DMSO-*d*_6_) δ 7.52 (s, 1H), 7.50 – 7.38 (m, 5H), 7.29 (t, *J* = 7.12 Hz, 2H), 7.24 (d, *J* = 7.17 Hz, 1H), 7.18 (d, *J* = 6.77 Hz, 2H), 6.02 (t, *J* = 5.47 Hz, 1H), 4.54 (d, *J* = 5.41 Hz, 2H); ^13^C NMR (101 MHz, DMSO-*d*_6_) δ 167.8, 158.1, 155.3, 138.1, 135.2, 134.8, 129.5, 129.4, 129.1, 128.9, 127.8, 127.6, 122.2, 112.8, 44.7; HRMS (ES+) m/z calc. for [C_19_H_14_ClN_3_S + H^+^]: 352.0670; found: 352.0663.

*4-(2-((2-chloro-5-phenylthieno[2,3-d]pyrimidin-4-yl)amino)ethyl)phenol (****5g***): Off white solid; Yield 32%; ^1^H NMR (850 MHz, DMSO-*d*_6_) δ 9.21 (s, 1H), 7.49 (s, 1H), 7.47 (d, *J* = 7.46 Hz, 1H), 7.43 (t, *J* = 7.52 Hz, 2H), 7.33 (d, *J* = 6.92 Hz, 2H), 6.81 (d, *J* = 8.41 Hz, 2H), 6.64 (d, *J* = 8.45 Hz, 2H), 5.57 (t, *J* = 5.35 Hz, 1H), 3.54 (q, *J* = 6.60 Hz, 2H), 2.62 (t, *J* = 6.72 Hz, 2H); ^13^C NMR (214 MHz, DMSO-*d*_6_) δ 170.4, 160.8, 158.9, 158.1, 137.9, 137.4, 132.5, 132.2, 131.8, 131.8, 131.7, 124.8, 118.4, 115.3, 45.5, 36.1; HRMS (ES+) m/z calc. for [C_20_H_16_ClN_3_OS + H^+^]: 382.0775; found: 382.0769.

*2-chloro-4-((4-hydroxybenzyl)amino)-5-phenylthieno[2,3-d]pyrimidin-6(5H)-one (***6b**): To a reaction vial containing compound **6a** (80 mg, 0.2 mmol, 1 equiv) in anhydrous DCM (6 mL) at 0 °C was added boron tribromide (2.5 equiv in 1.5 mL DCM) dropwise under nitrogen atmosphere over 5 min. The reaction was continued to stir at 0 °C until the disappearance of starting material (4h per TLC). The reaction content were then poured slowly into the ice-water mixture (50 mL) and extracted with DCM (20 mL) twice. The organic layer was washed with Sat. bicarbonate if necessary, finally the organic layer was dried on sodium sulfate, evaporated to yield crude residue which was purified through flash chromatography by gradient elution (Ethyl acetate : Hexane). White solid; Yield 51%; ^1^H NMR (400 MHz, DMSO-*d*_6_) δ 9.23 (s, 1H), 7.45 (t, *J* = 5.97 Hz, 1H), 7.41 – 7.29 (m, 3H), 7.20 (dd, *J* = 6.70, 2.77 Hz, 2H), 6.76 (d, *J* = 8.35 Hz, 2H), 6.56 (d, *J* = 8.48 Hz, 2H), 5.28 (s, 1H), 4.43 (dd, *J* = 15.05, 6.50 Hz, 1H), 4.24 (dd, *J* = 15.18, 5.37 Hz, 1H); ^13^C NMR (214 MHz, DMSO-*d*_6_) δ 202.1, 169.5, 161.6, 159.9, 159.4, 136.0, 132.1, 131.6, 131.5, 131.2, 131.2, 118.0, 112.5, 62.4, 46.1; HRMS (ES+) m/z calc. for [C_19_H_14_ClN_3_O_2_S + H^+^]: 384.0568; found: 384.0562.

### ERα LiSA

HepG2 cells were maintained in Basal Medium Eagles supplemented with 8% fetal bovine serum (FBS) and plated (2.5XE06 cells/10mL) in T75 flasks for 24 hours prior to transfection. Cells were transfected with VP16-ERα, 5XGalLuc3, pM-alpha II, and Renilla luciferase at a 1:30:1:5 ratio using Lipofectin (Thermo Fisher) according to manufacturer’s protocol. Following 24 h transfection, cells were plated in 384-well plates and treated for 24 hours with 1.0 μM of 4OHT, FULV, BZE, RAL, RAD, or 10 μM of the phenol library compounds. Dual luciferase assays were performed from cell extracts. Renilla luciferase served as control for cellular toxicity. Data is presented as light units.

### ERE reporter gene assay

HepG2 cells were plated in T75 flasks (1.5XE06 cells/10mL) and sub-cultured for 24 h. Cells were transfected with pRST7-ERα, Renilla luciferase, and a 7X-ERE-TK-luc reporter gene at a 1:4:25 ratio using Lipofectin transfection reagent. Following 24 h transfection, cells were treated with E2 (1.0E-12–1.0E-06 M), FULV, 4OHT, or compound **2a** (1.0E-11–1.0E-05 M) for 24 h. Cells were lysed and extracts were assayed for Firefly and Renilla luciferase activity using dual luciferase reagent (DLR). Data is presented as light units and error bars represent S.D. of triplicate points.

### ER whole cell competition binding assay

1.0E05 MCF7 cells were plated in DMEM/F12 with 8% charcoal-stripped fetal bovine serum in 24-well plates for 24 h. Following incubation, cells were treated with E2, compound **2a**, or compound **6b** (1.0E-06–1.0E-10 M) in the presence of 0.1 nM ^3^H-E2. To determine background levels of radioactivity, control wells were treated with 500X cold E2 (50 nM). After 2 h incubation, cells were lysed using 200 μL lysis buffer (2% SDS, 10% Glycerol, 10 mM Tris-HCl pH 6.8); and then volumes were increased to 500 μL using 10 mM Tris-HCl (pH 8.0). 300 μl of the lysates were added to 3 mL of Cytoscint (MP Biomedicals) and analyzed by scintillation counting (Beckman LS 6000SC).

### Protein Expression and Purification

A His_6_-TEV-tagged ERα LBD (305–555) with a Y537S mutation in pET21a was expressed in *E*.*coli* BL21(DE3) using the previously reported methods^17^. Cells were grown at 37 °C with shaking in LB broth, supplemented with ampicillin, until they reached mid-log phase growth (OD_600_ = 0.6). The temperature was reduced to 16 °C and protein expression was induced with 0.4 M IPTG for 16 h. Cells were harvested by centrifugation at 4,000 *xg* for 15 min then resuspended with 20 mL/g cell paste with 50 mM HEPES pH 8.0, 500 mM NaCl, 0.5 mM TCEP, 20 mM imidazole pH 8.0, 5% glycerol, and EDTA-free PIC tabs (Pierce). Cells were lysed by sonication and cellular debris was cleared by centrifugation at 18,000 *xg* for 45 min. A BioRad NGC Quest FPLC was used for protein purification. The supernatant containing the His_6_-TEV-ERα was captured using a BioRad Nuvia IMAC column, washed with lysis buffer until baseline was reached, and eluted with 50 mM HEPES pH 8.0, 500 mM NaCl, 0.5 mM TCEP, 500 mM imidazole pH 8.0, and 5% glycerol. 1: 5,000 mol:mol His-TEV protease was added to the protein and the solution was dialyzed against 4 L of 50 mM HEPES pH 8.0, 500 mM NaCl, 0.5 mM TCEP, pH 8.0, 5% glycerol overnight at 4°C with stirring for 16 hours. The mixture was placed back over the IMAC column to remove the His-tag and His-TEV protease. ERα LBD was purified a final time using a GE Superdex 16/600 size exclusion column. A single peak corresponding to approximately 50 kDa MW corresponding to the ERα LBD dimer was concentrated to 10-15 mg/mL, flash frozen, and stored at -80°C.

### Crystallization, X-Ray Data Collection, and Structure Solution

Each small molecule ligand was added to 10 mg/mL ERα LBD Y537S at 1 mM alongside GRIP peptide at 3 mM overnight at 4°C. Hanging drop vapor diffusion was used to crystallize the protein using Hampton VDX plates. Clear rectangular crystals emerged after 16 hours at room temperature in 15–25% PEG 3,350, 100 mM MgCl_2_, 100 mM Tris pH 8.0 at 5 mg/mL. Paratone-N was used as a the cryoprotectant. X-ray diffraction data were collected at the Structural Biology Center 19-BM beamline at the Advanced Photon Source, Argonne National Laboratories at 0.97 Å. Data were indexed and scaled using HKL 3,000.^28^ Phenix^29^ was used for molecular replacement using PDB 6CBZ as the input model.^30^ The structures were refined using iterative rounds of Phenix Refine and manual inspection/structure editing using Coot. Significant difference density was observed in the ligand binding pockets with one round of structure refinement that resolved after placing the compounds in the ligand binding pocket and further refinement. X-ray crystal structure collection and refinement statistics are recorded in SI Table 3. The coordinates and map were deposited in the protein databank (PDB) with accession codes 7RKE and 7T2X.

## ASSOCIATED CONTENT

### Supporting Information

SI Table 1 – Primary hits from ERα LiSA screen

SI Table 2 – Commercial thienopyrimidine analogs

SI Table 3 – X-ray refinement data

SI Table 4 – Kinase Tm screen

SI Figure 1 – Electron density maps of **2a** and **6b**

SI Figure 2 – **6b/6b’** keto/enol NMR ratios

SI Scheme 1 – Gewald reaction mechanism

NMR spectra of all final compounds **2a**–**g, 5a**–**g**, and **6a**–**b**

## ACKNOWLEDGEMENTS

The SGC is a registered charity (number 1097737) that receives funds from AbbVie, Bayer Pharma AG, Boehringer Ingelheim, Canada Foundation for Innovation, Eshelman Institute for Innovation, Genome Canada, Genentech, Innovative Medicines Initiative (EU/EFPIA) [ULTRA-DD grant no. 115766], Janssen, Merck KGaA Darmstadt Germany, MSD, Novartis Pharma AG, Ontario Ministry of Economic Development and Innovation, Pfizer, São Paulo Research Foundation-FAPESP, Takeda, and Wellcome [106169/ZZ14/Z]. This work was supported by the Department of Defense through the FY 17 BRCP Innovator Award under Award No. BC170954. Opinions, interpretations, conclusions, and recommendations are those of the authors and are not necessarily endorsed by the Department of Defense. Research reported in this publication was supported in part by the NC Biotech Center Institutional Support Grant 2018-IDG-1030. Funding from Loyola University Chicago Startup Funds and Susan G. Komen CCR 19608597. Results shown in this report are derived from work performed at Argonne National Laboratory (ANL), Structural Biology Center (SBC) at the Advanced Photon Source (APS), under U.S. Department of Energy, Office of Biological and Environmental Research contract DE-AC02-06CH11257. We thank Dr. Stefan Knapp (SGC-Frankfurt) for the kinase selectivity data.

