## Supporting Information for "A new chemotype of chemically tractable nonsteroidal estrogens based on a thieno[2,3-*d*]pyrimidine core"

**Table of Contents**

**P2–9**	SI Table 1 – Primary hits from ERα LiSA screen

**P10–12**	SI Table 2 – Commercial thienopyrimidine analogs

**P13**	SI Table 3 – X-ray crystallographic refinement data

**P14**	SI Table 4 – Kinase selectivity screen of **2a**

**P15**	SI Figure 1 – Electron density maps of **2a** and **6b**

**P16**	SI Figure 2 – **6b/6b’** keto/enol NMR ratios

**P17**	SI Scheme 1 – Gewald reaction mechanism

**P18–33**	NMR spectra of all final compounds **2a**–**g**, **5a**–**g**, and **6a**–**b’**

SI Table 1 – Primary hits from ERα LiSA screen

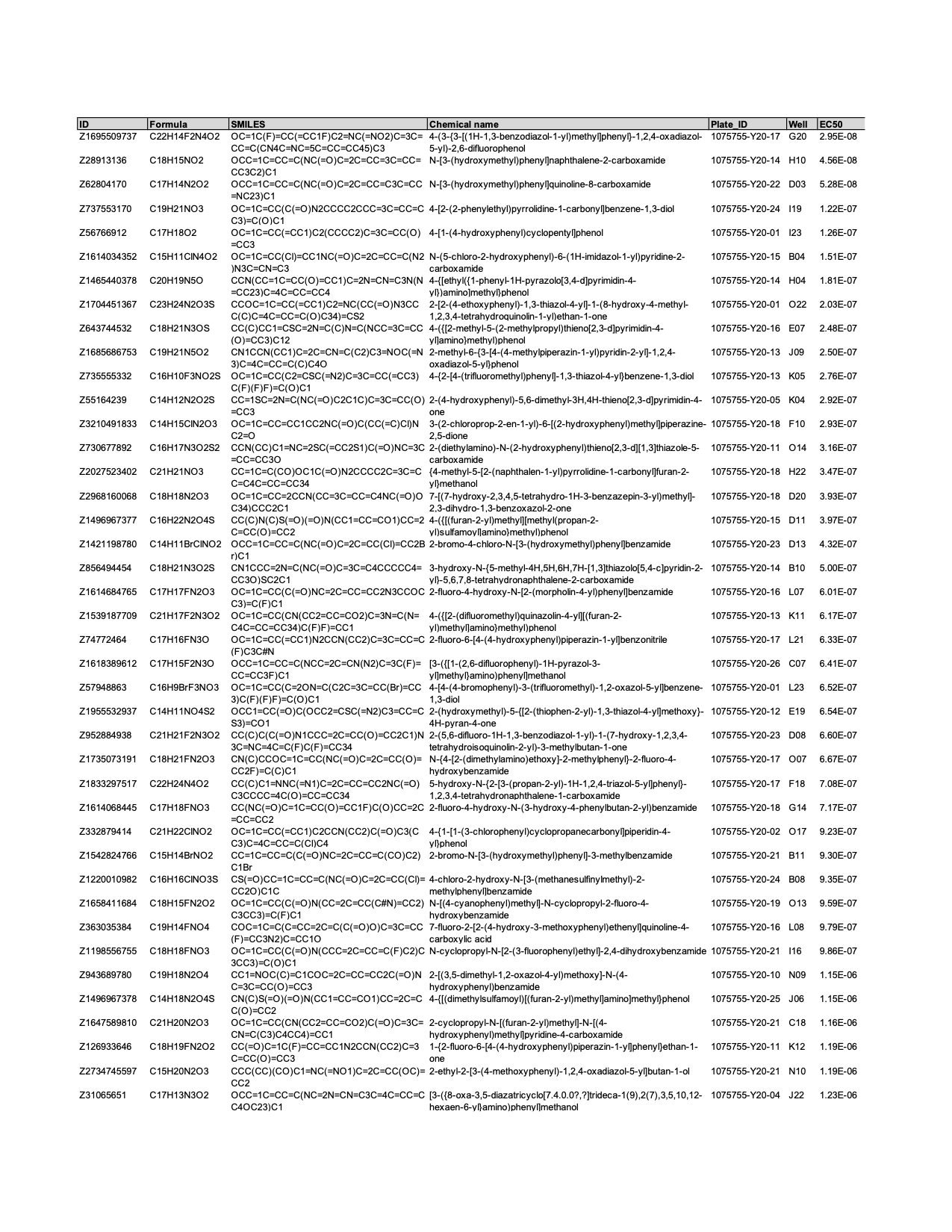

SI Table 2 – Commercial thienopyrimidine analogs

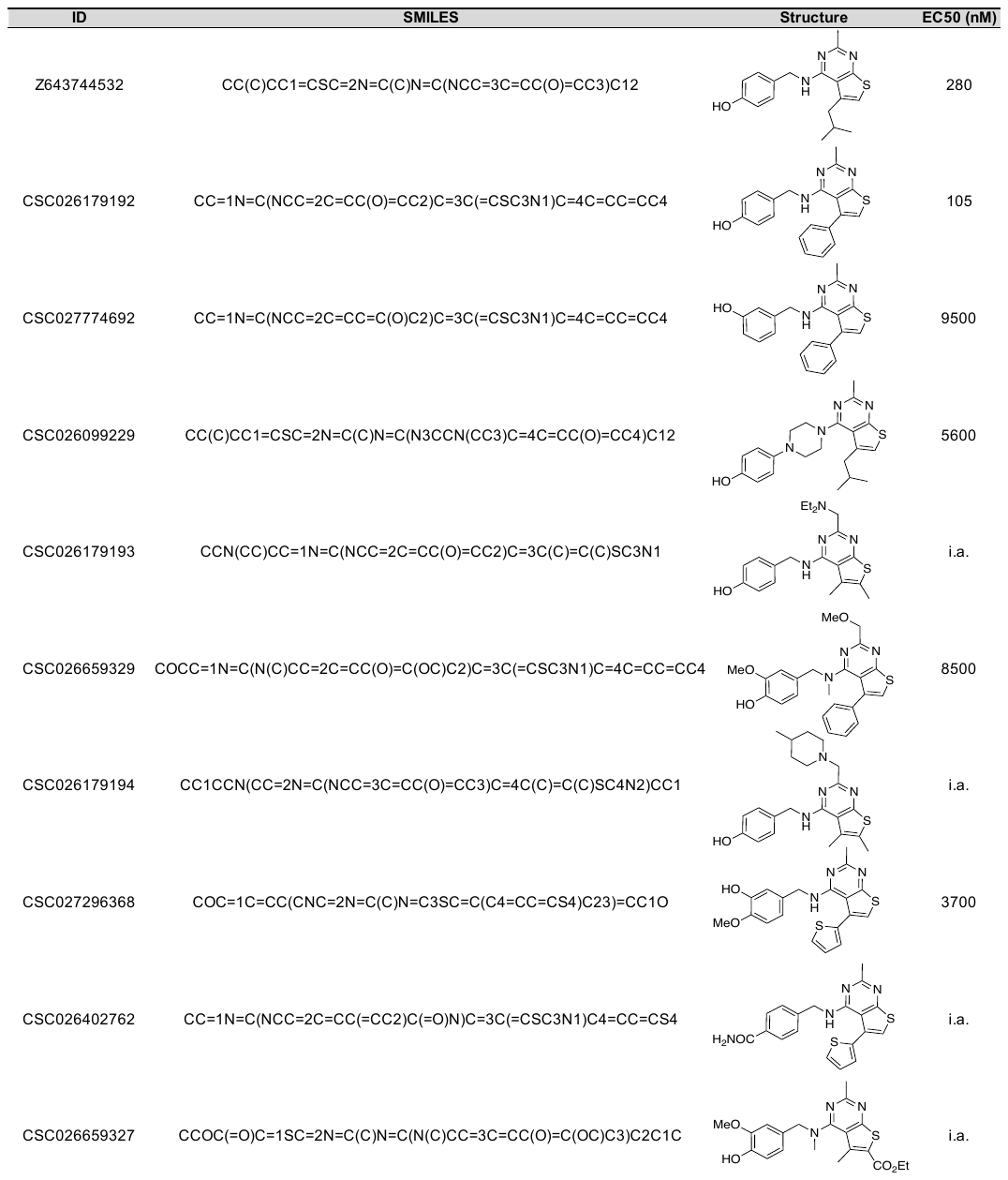

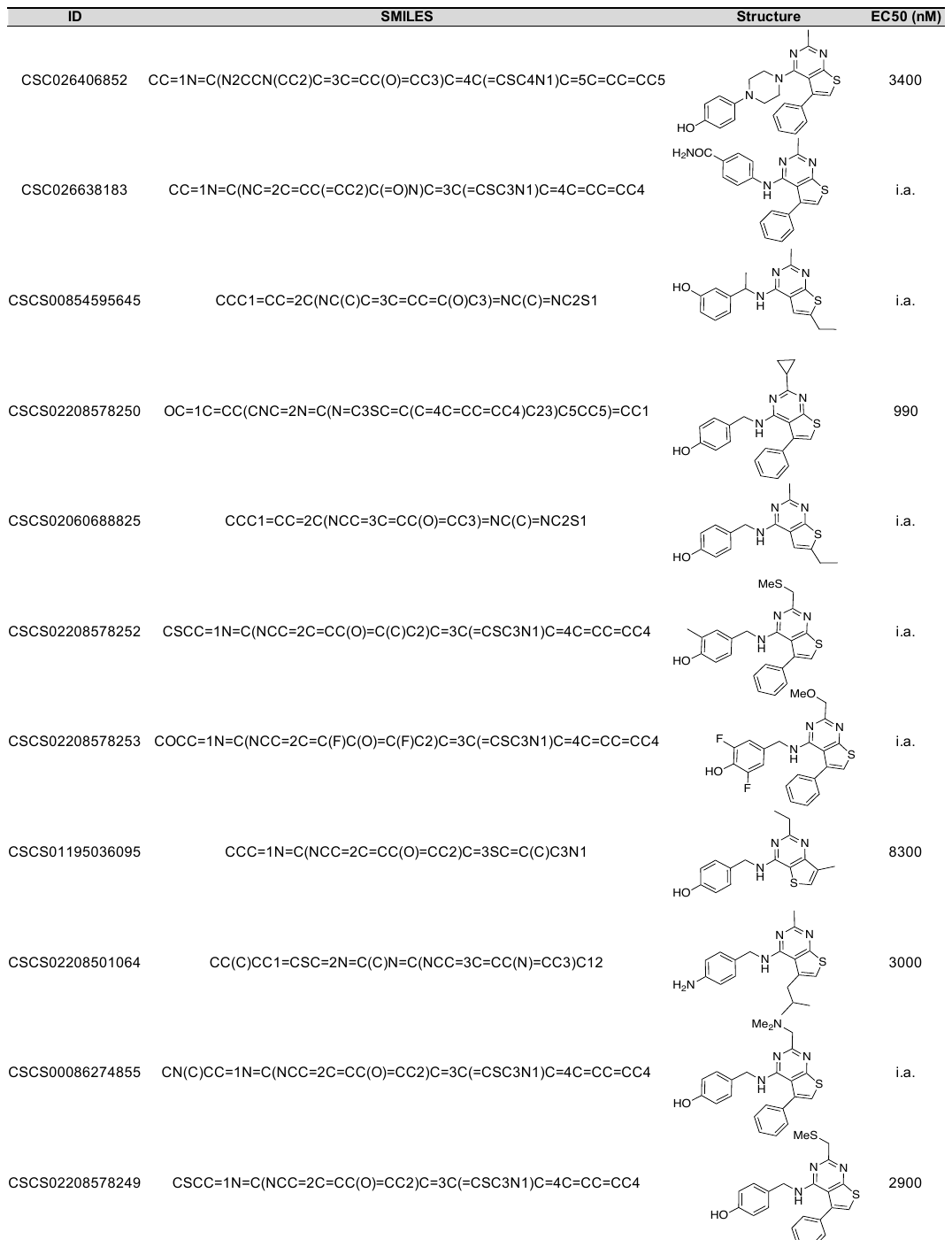

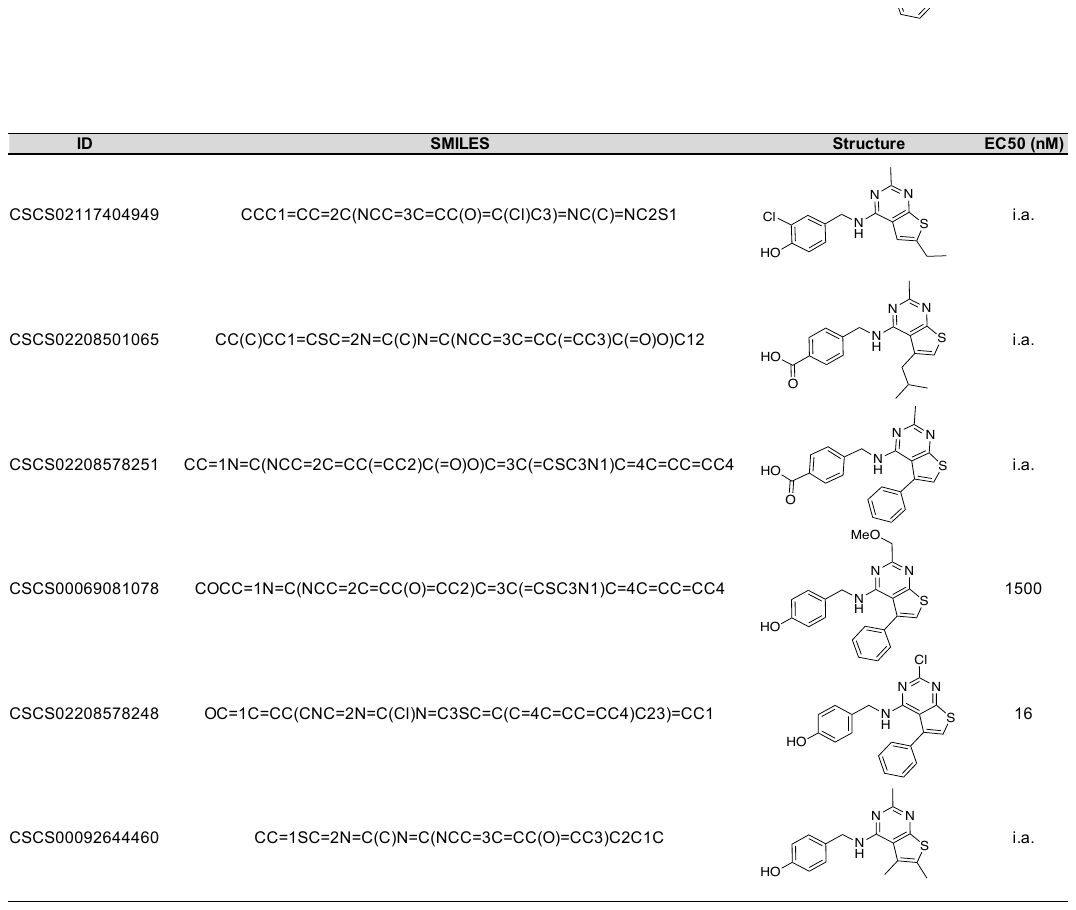

SI Table 3 – X-ray crystallographic refinement data

|  | **ERα LBD Y537S** | **ERα LBD Y537S** |
| --- | --- | --- |
|  | 4-(((2-chloro-5-phenylthieno[2,3-d]pyrimidin-4-yl)amino)methyl)phenol (**2a**) | 2-chloro-4-((4-hydroxybenzyl)amino)-5-phenylthieno[2,3-d]pyrimidin-6-ol (**6b**) |
| PDB ID | 7RKE | 7T2X |
| **Data Collection** | |  |
| Space Group | P1211 | P1211 |
| a, b, c (Å) | 56.03, 83.94, 58.66 | 56.13, 84.04, 58.91 |
| α, β, γ (°) | 90.00, 108.66, 90.00 | 90.00, 109.47, 90.00 |
| Resolution Range (Å) | 33.49-1.55 | 46.78-2.60 |
| Number of Reflections (Highest Resolution) | 60,986 (3,904) | 38,882 (1,702) |
| Completeness (Highest Resolution) | 90.3 (95.6) | 97.8 (86.8) |
| Redundancy | 3.6 | 3.6 |
| CC^1/2^ (Highest Resolution) | 1.000 (0.592) | 0.997 (0.626) |
| R_work_/R_free_ | 23.6/27.3 | 21.8/27.7 |
| No. Atoms | 4,137 | 3,595 |
| Water Atoms | 446 | 51 |
| Ligand Atoms | 50 | 52 |
| Bond Lengths (Å) | 0.014 | 0.004 |
| Bond Angles (°) | 1.15 | 0.59 |
| Preferred Number (%) | 99.11 | 98.62 |
| Additional Allowed (%) | 0.89 | 1.38 |
| Outliers (%) | 0 | 0 |

SI Table 4 – Kinase selectivity screen of **2a**

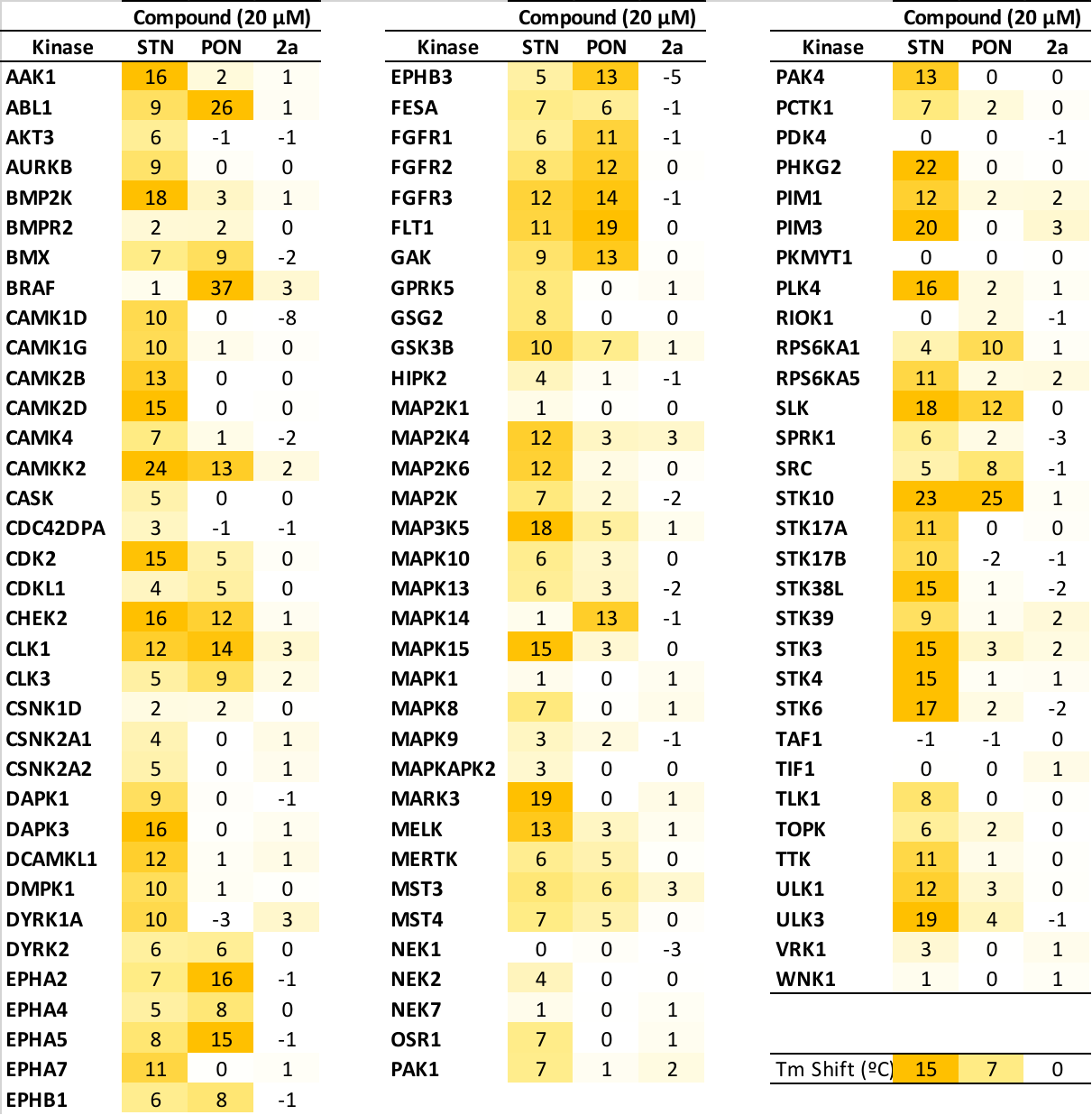

Thienopyrimidine **2a** did not stabilize the melting temperature (>2 °C or >30% of control) of 100 human protein kinases. Staurosporine (STN) and ponatinib (PON) were used as positive controls. All compounds were tested at a concentration of 20 μM. Data are shaded by the degree of thermal shift as shown in the legend.

SI Figure 1 – Electron density maps of **2a** and **6b**

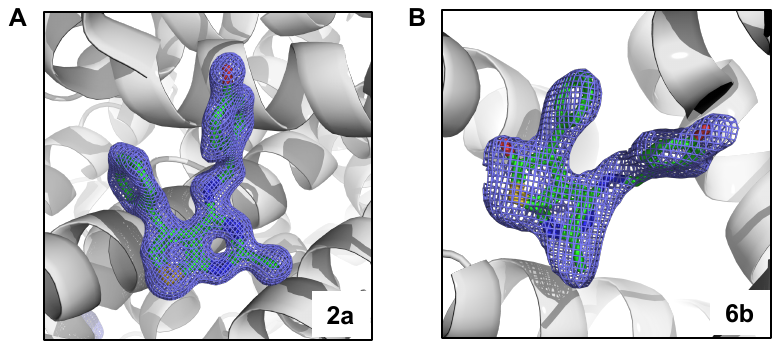

2mFo-DFc difference maps of **2a** (A) and **6b** (B) contoured to 2 σ. Electron density maps are shown as blue mesh, ligands are shown as sticks.

SI Figure 2 – **6b/6b’** keto/enol NMR ratios

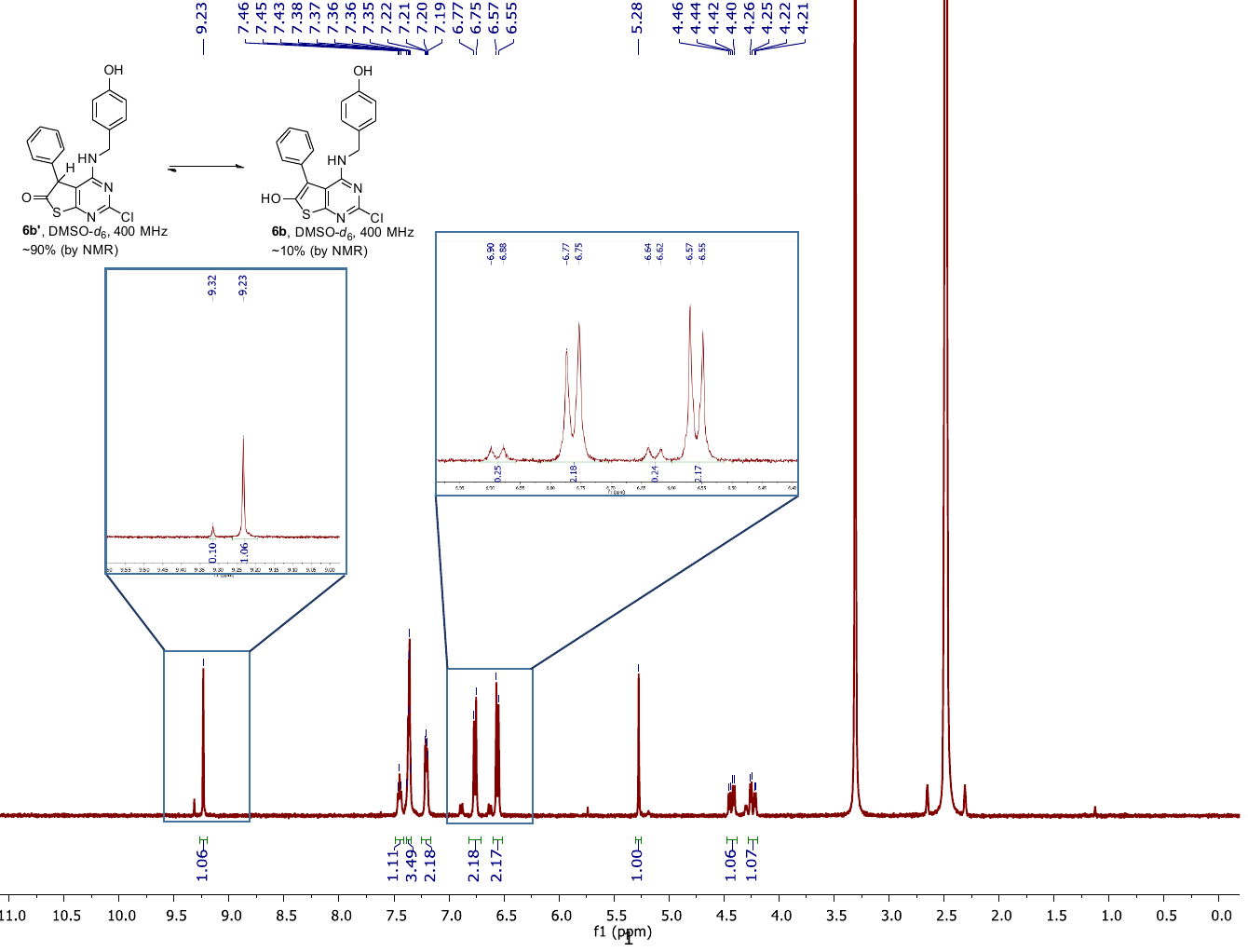

SI Scheme 1 – Gewald reaction mechanism

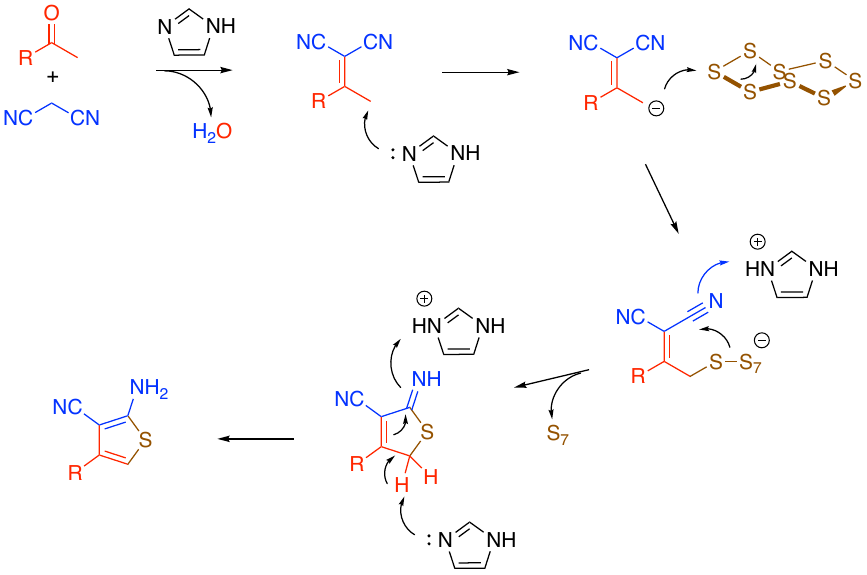

Proposed mechanism of the imidazole-catalyzed Gewald reaction showing the participation of a base that can both donate and accept a proton at multiple steps in the reaction sequence.

NMR spectra of all final compounds **2a**–**g**, **5a**–**g**, and **6a**–**b’**

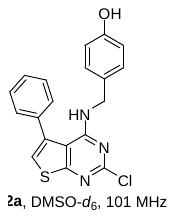

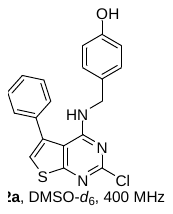

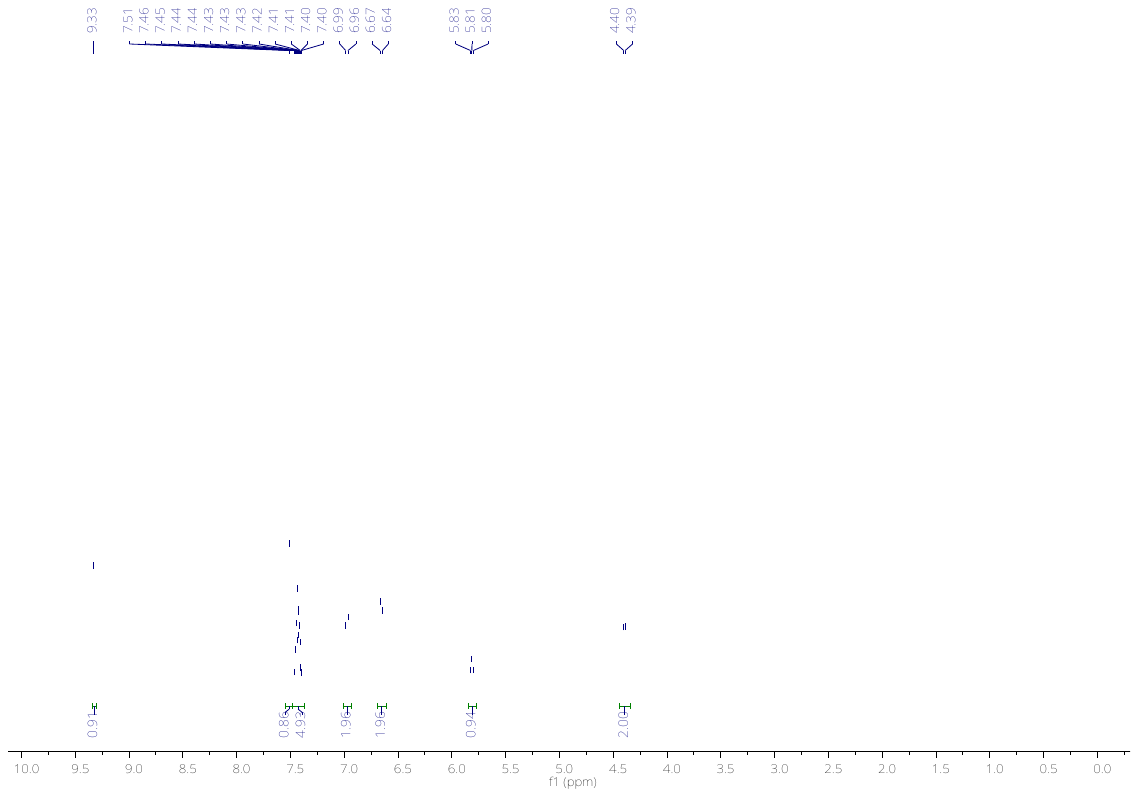

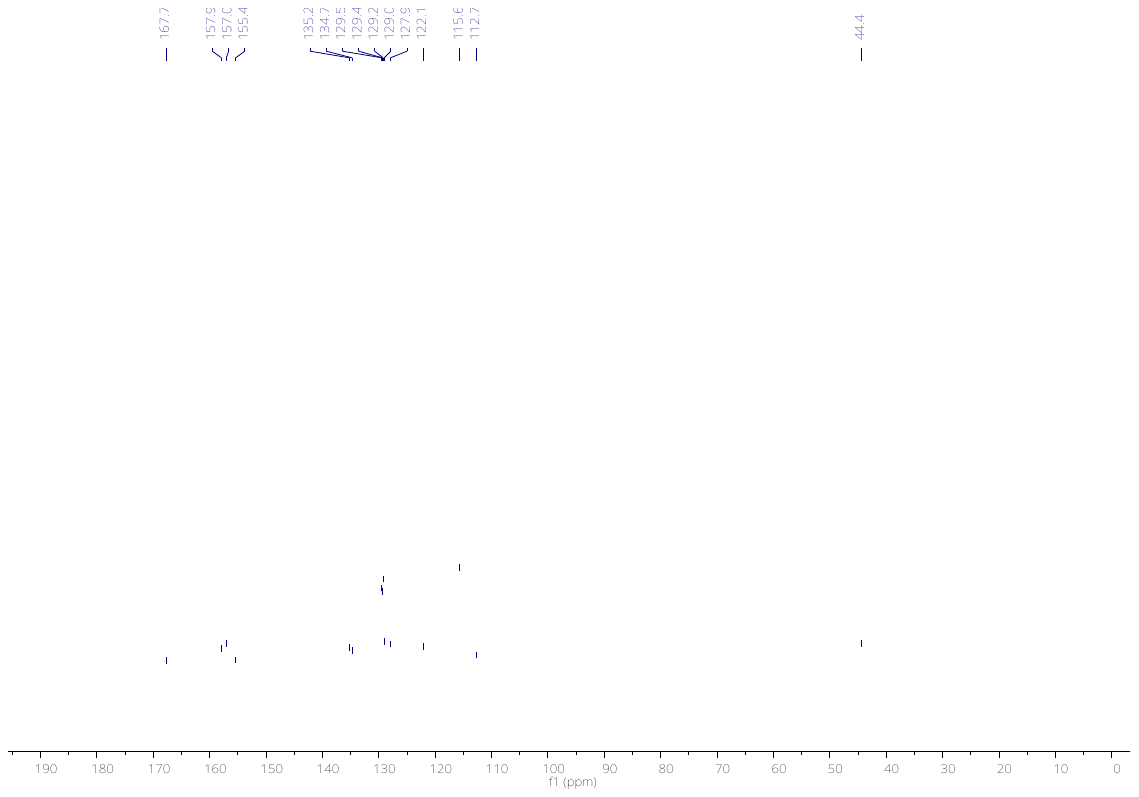

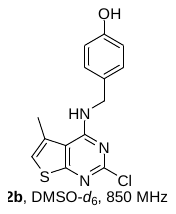

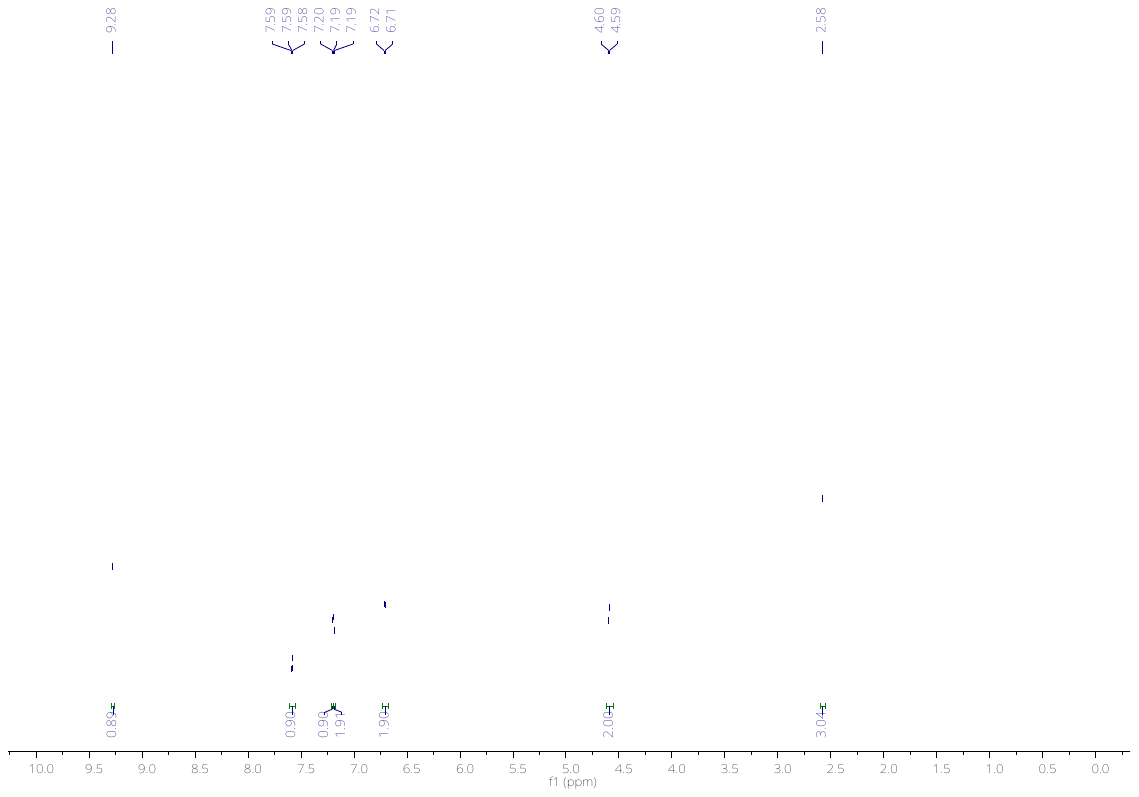

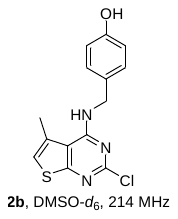

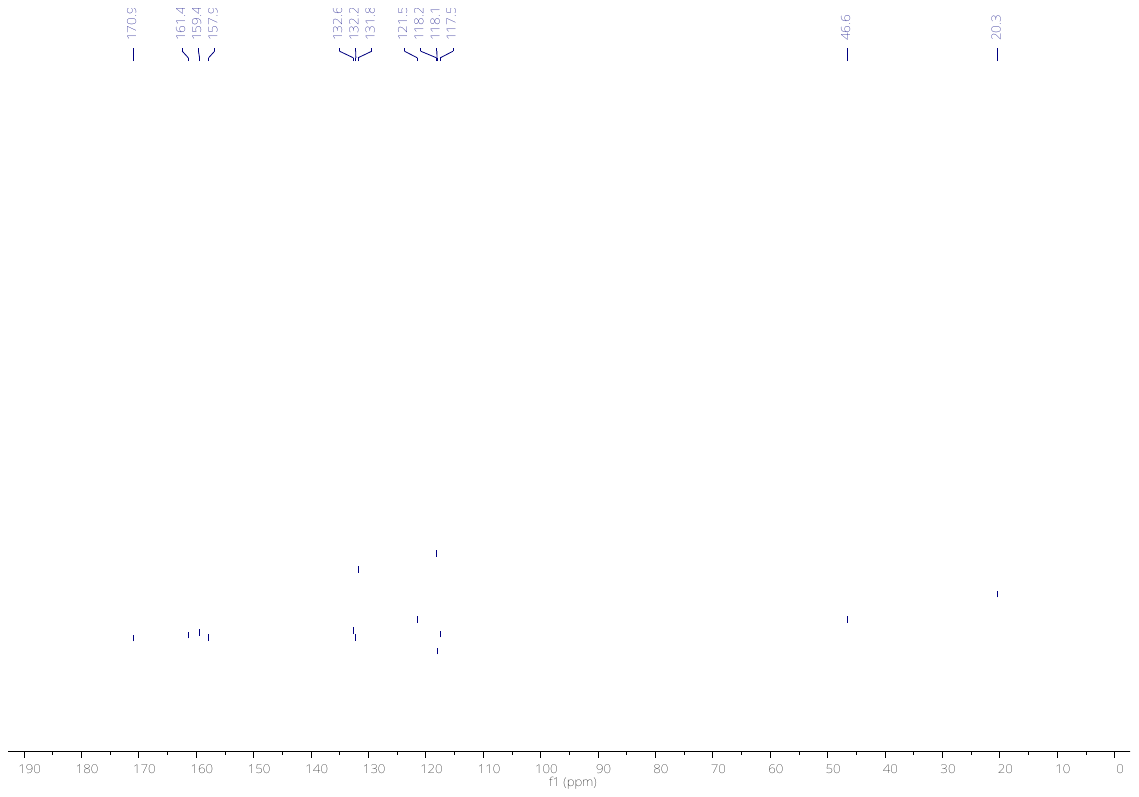

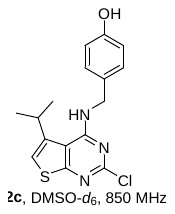

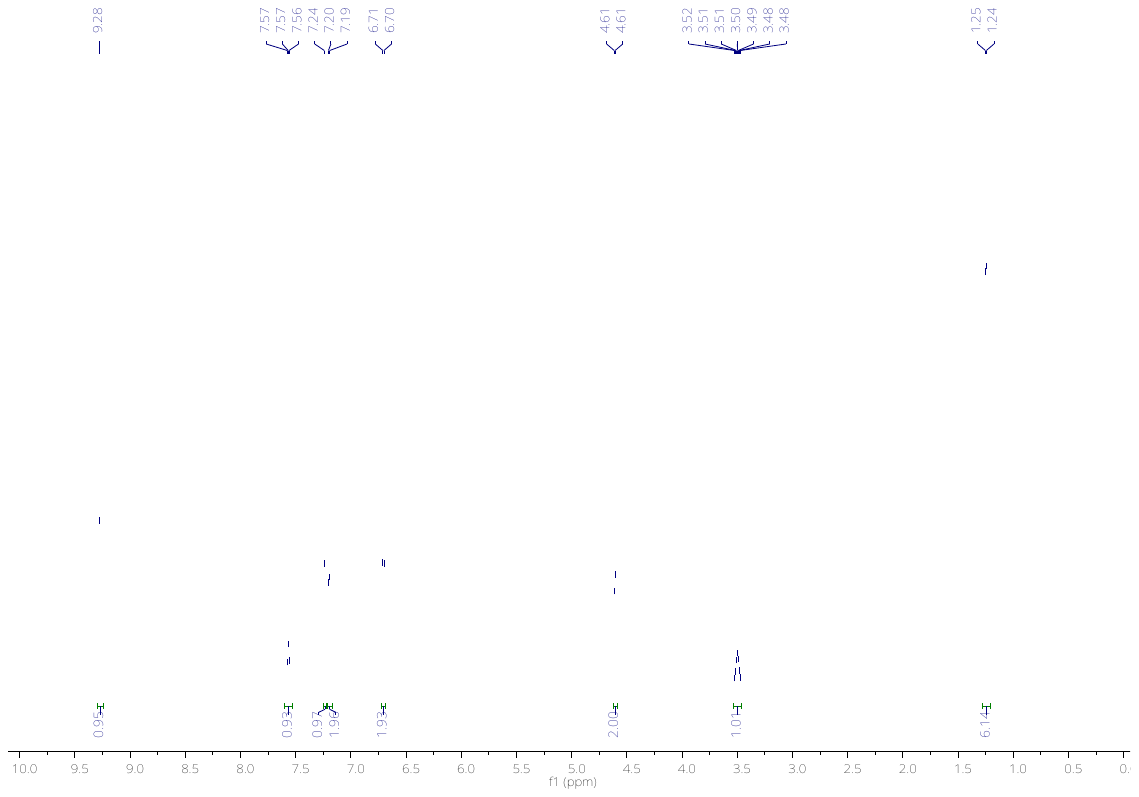

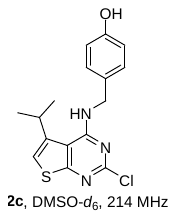

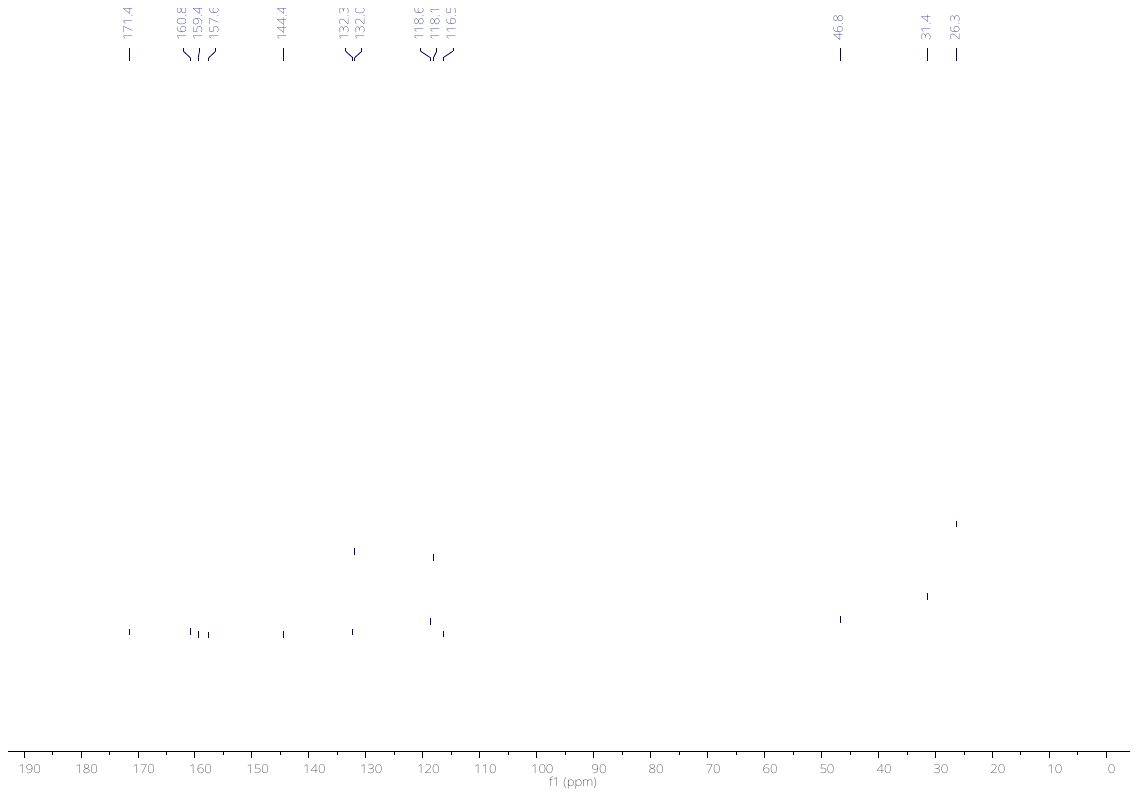

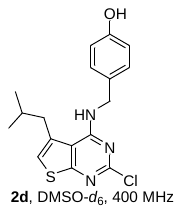

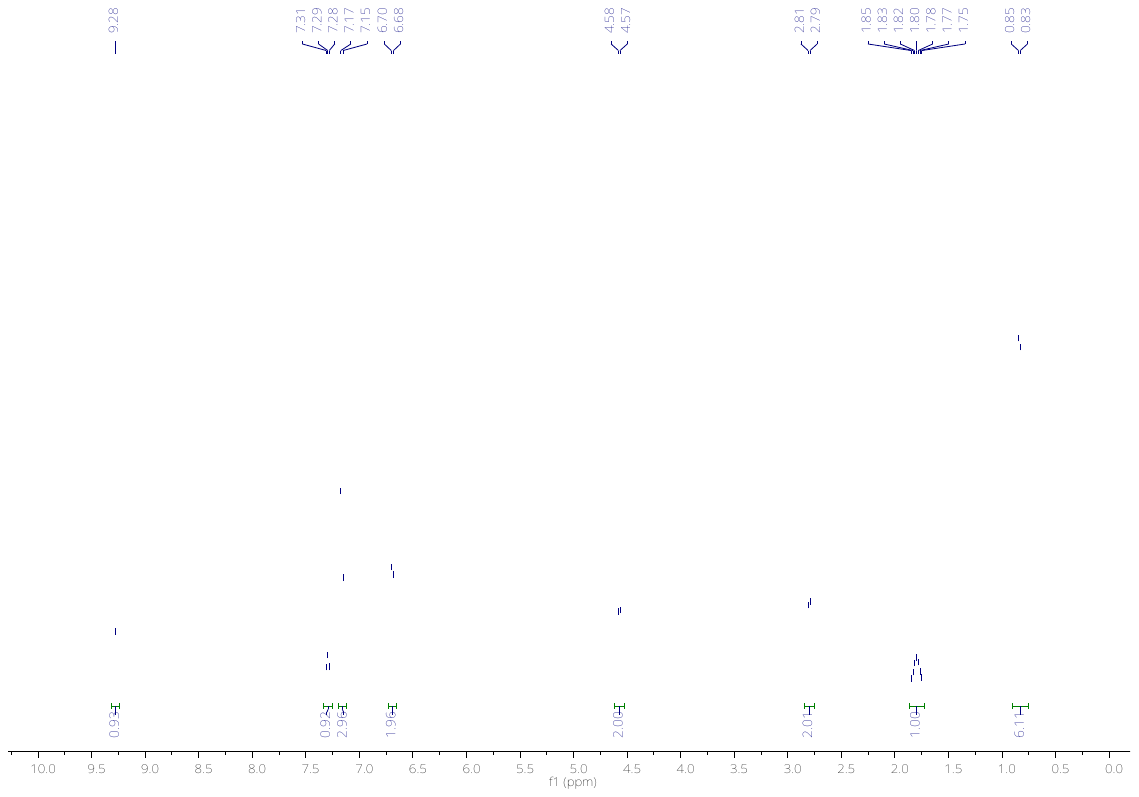

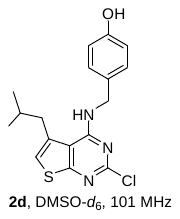

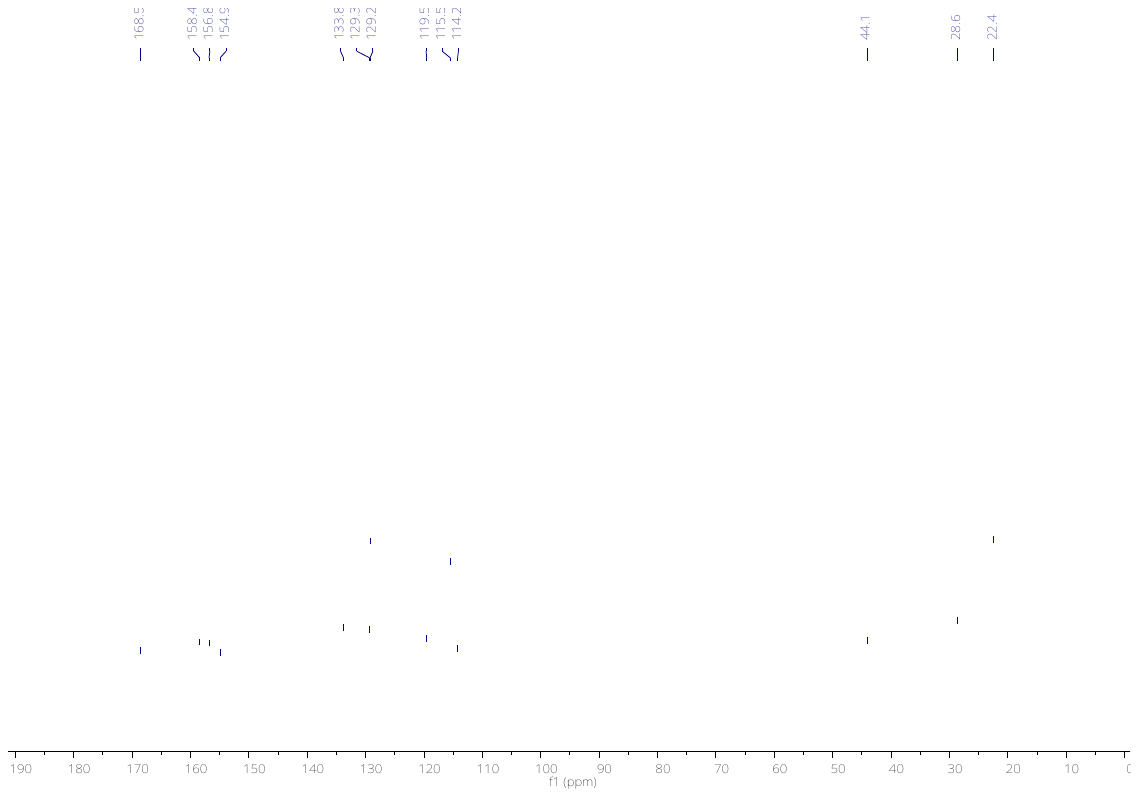

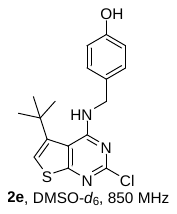

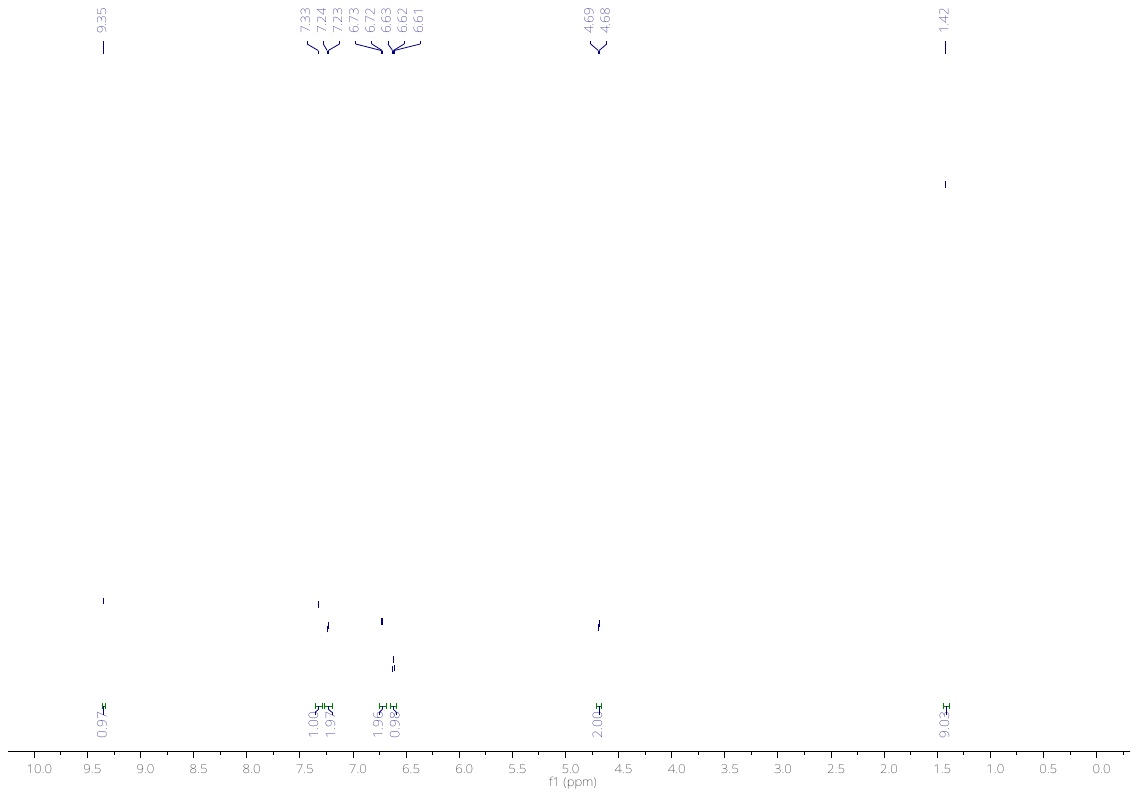

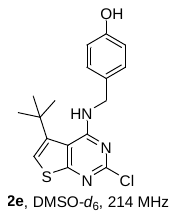

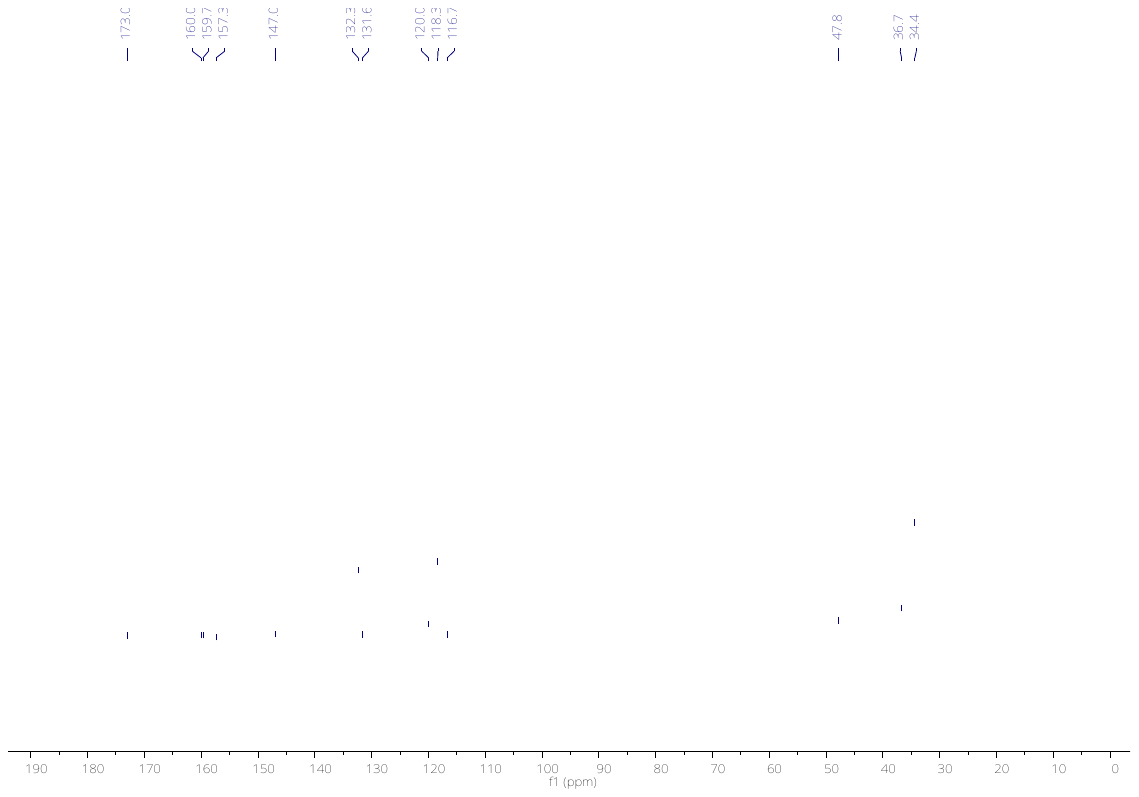

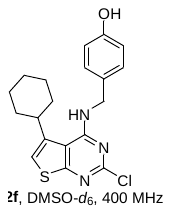

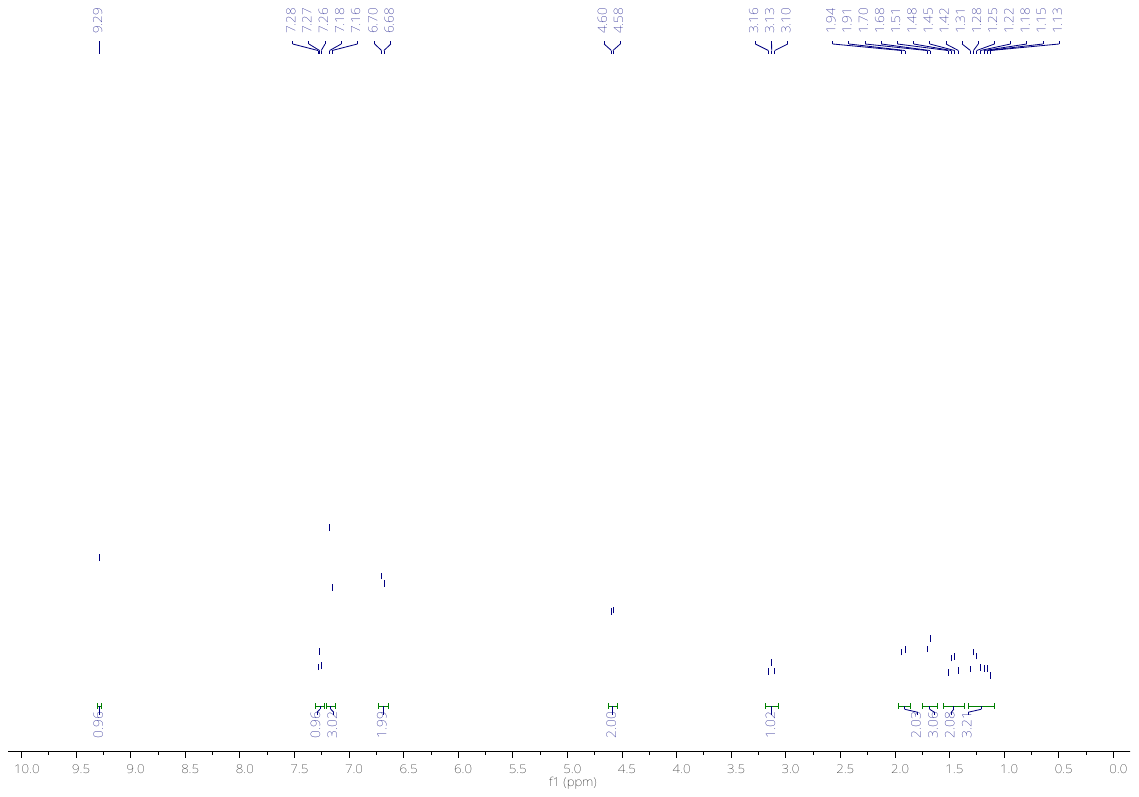

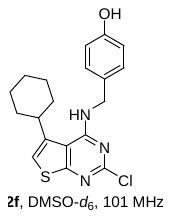

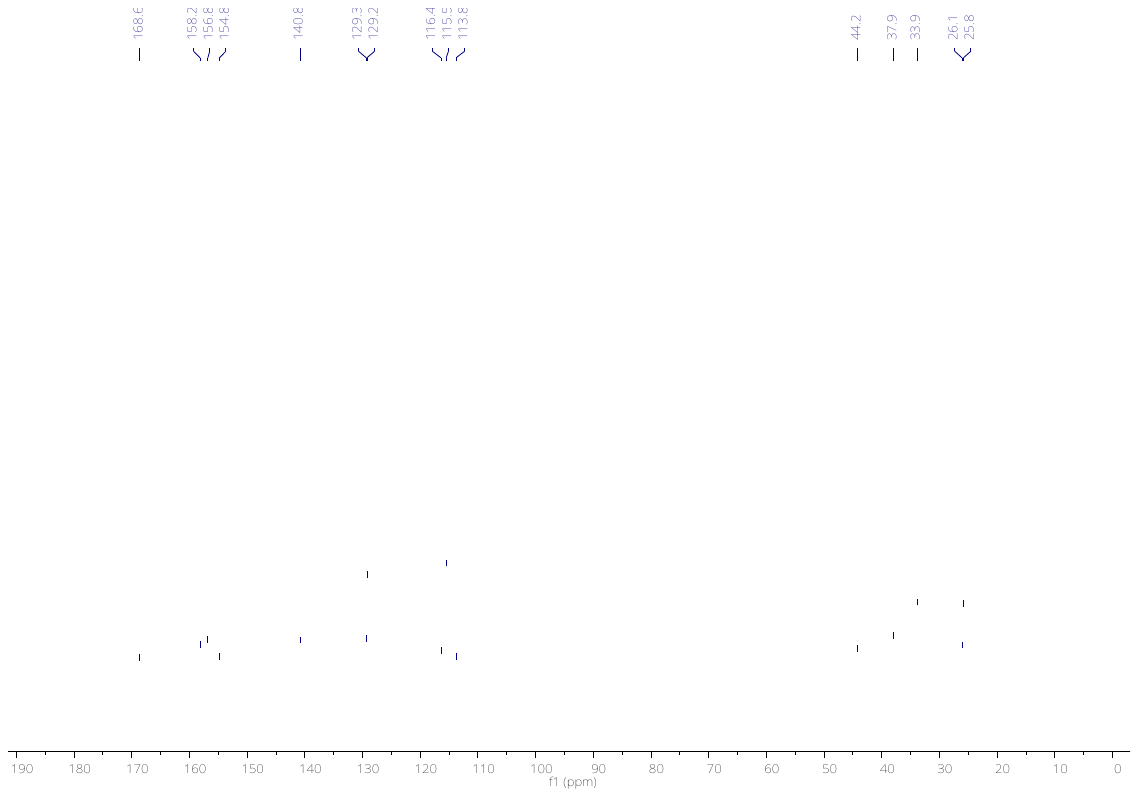

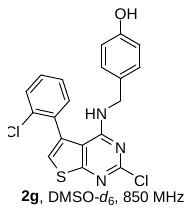

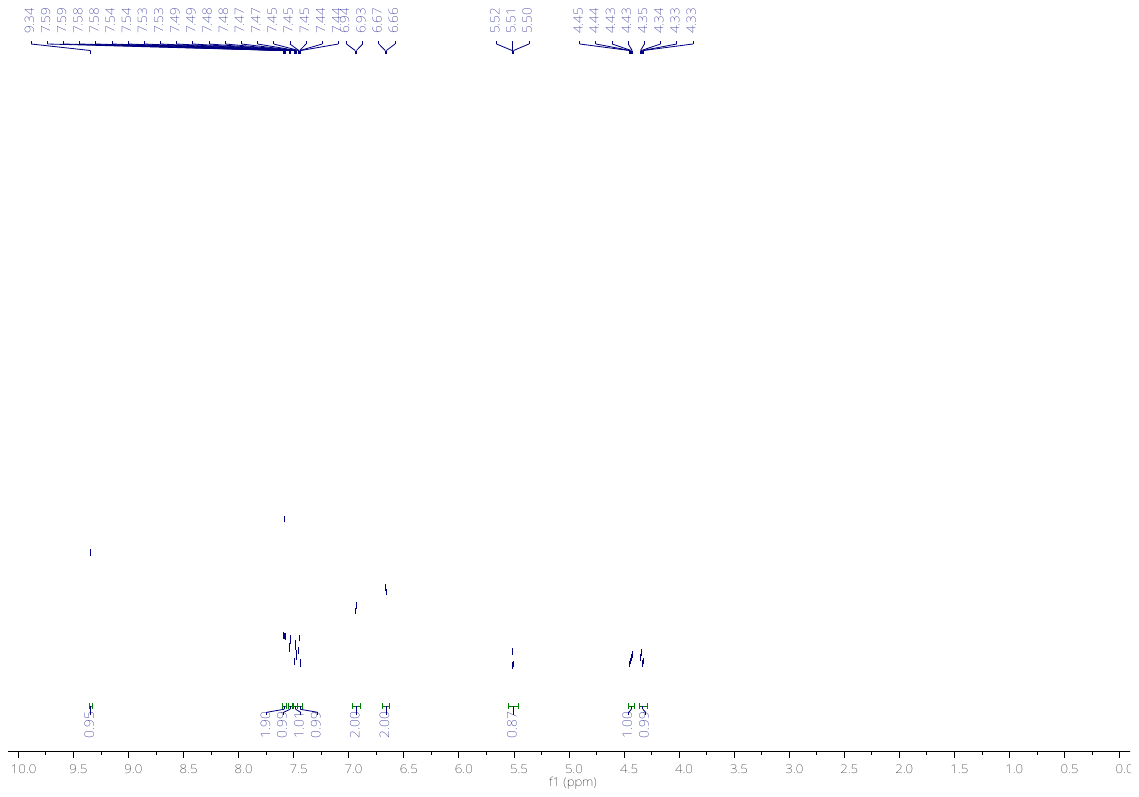

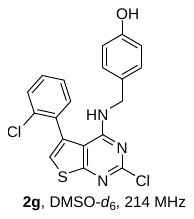

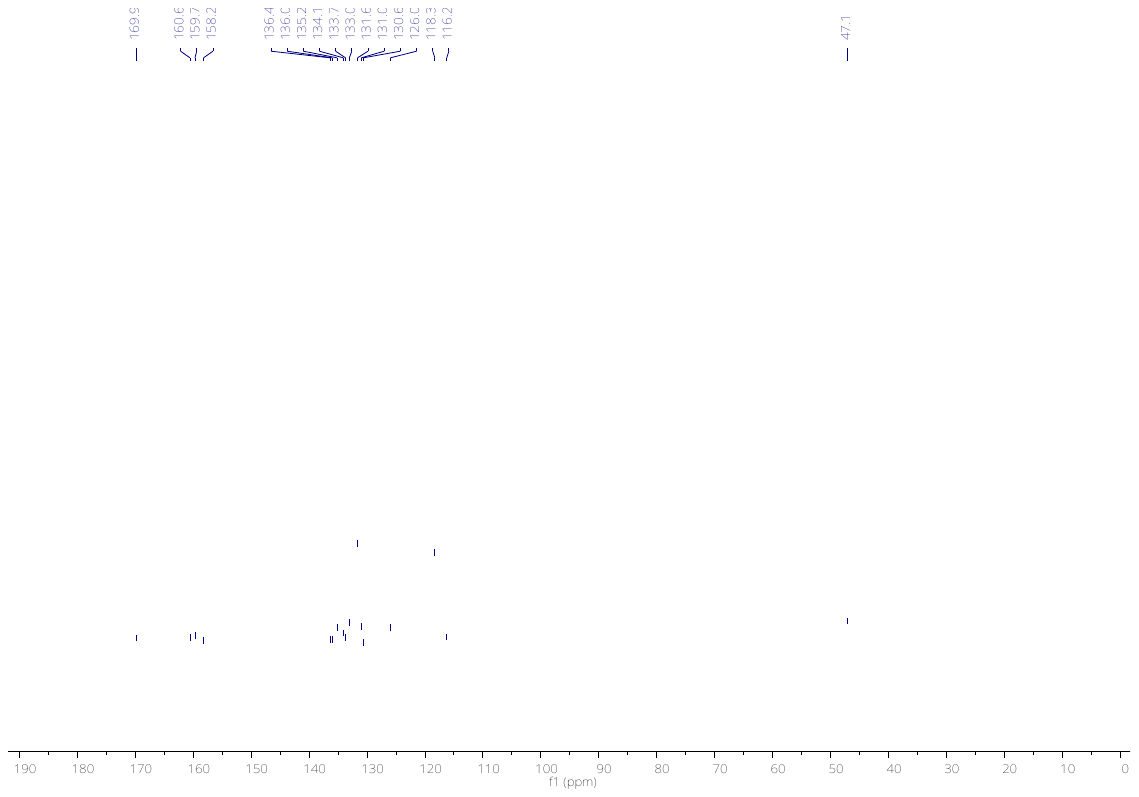

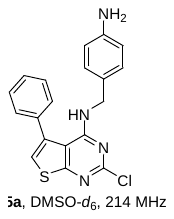

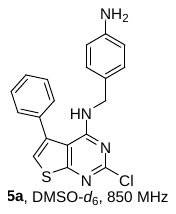

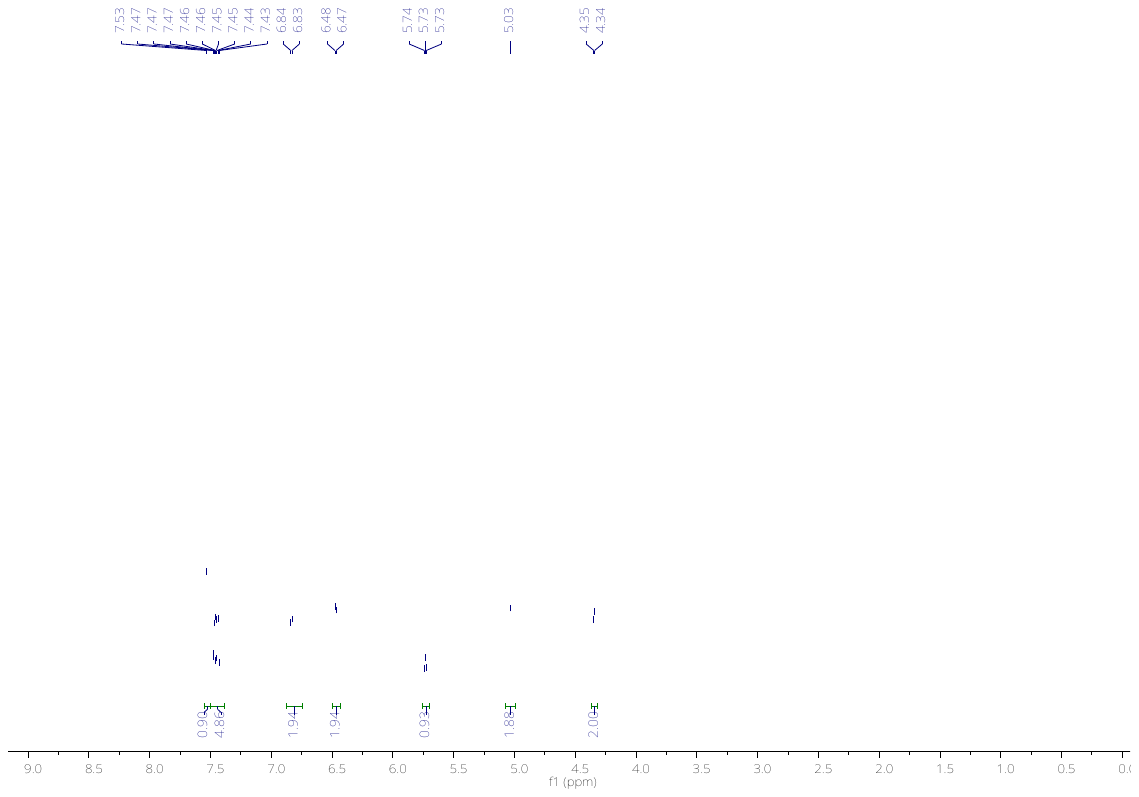

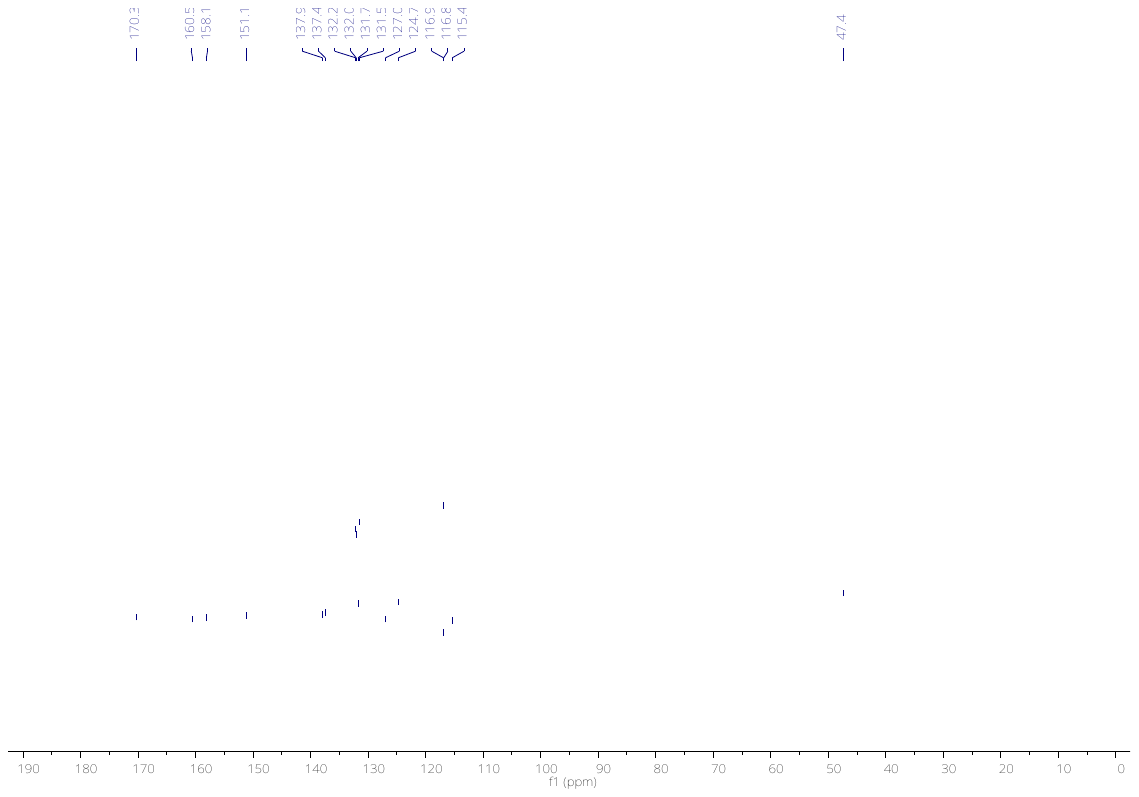

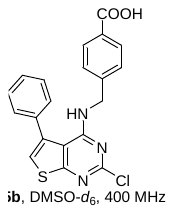

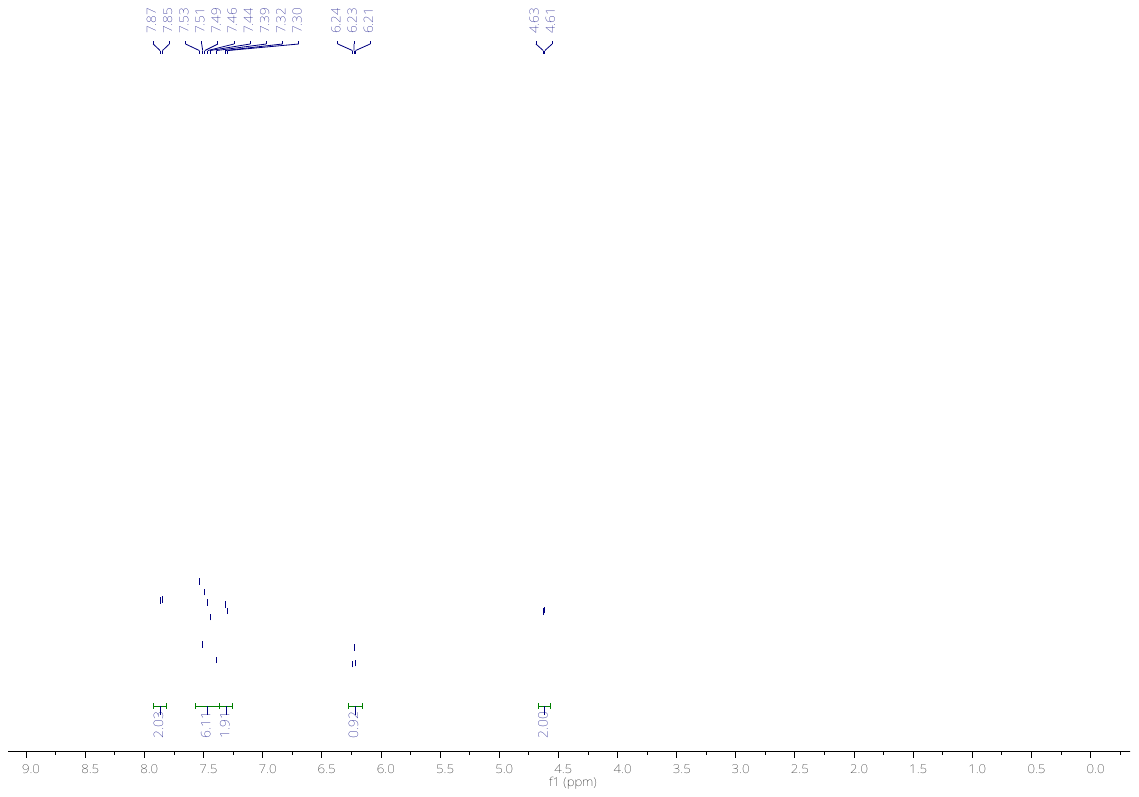

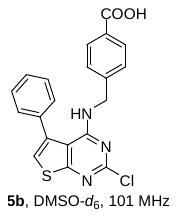

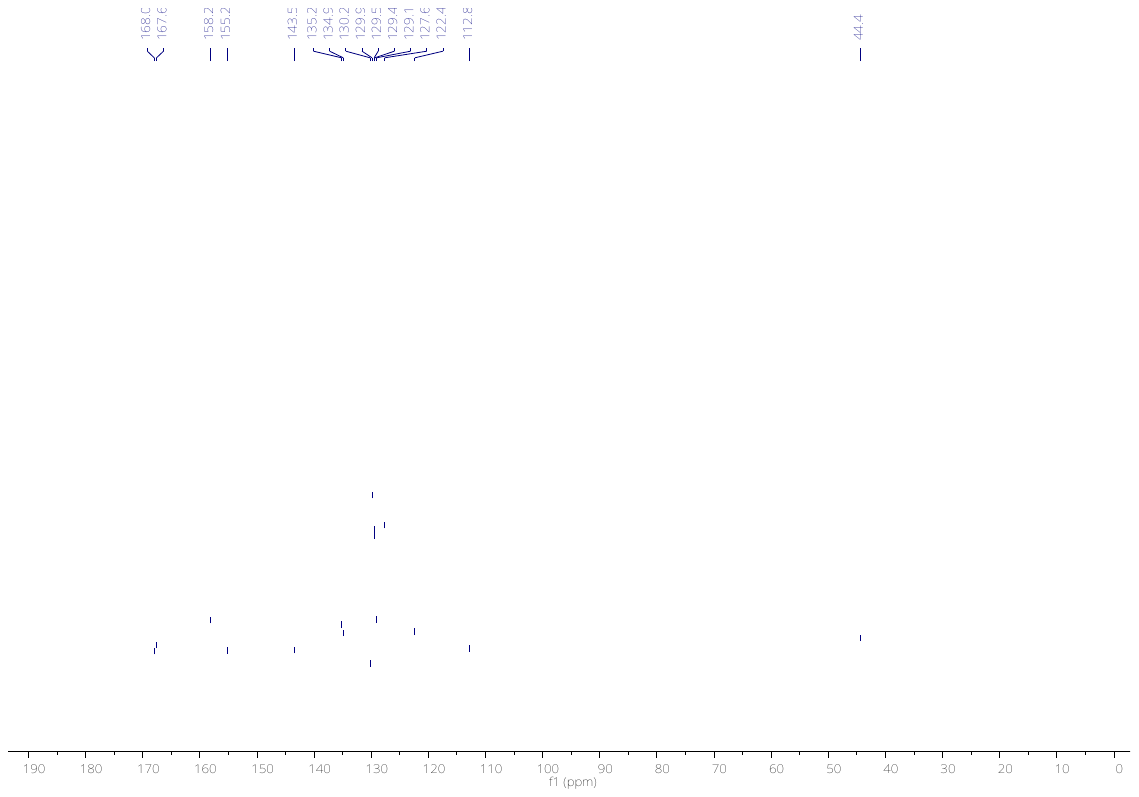

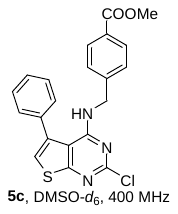

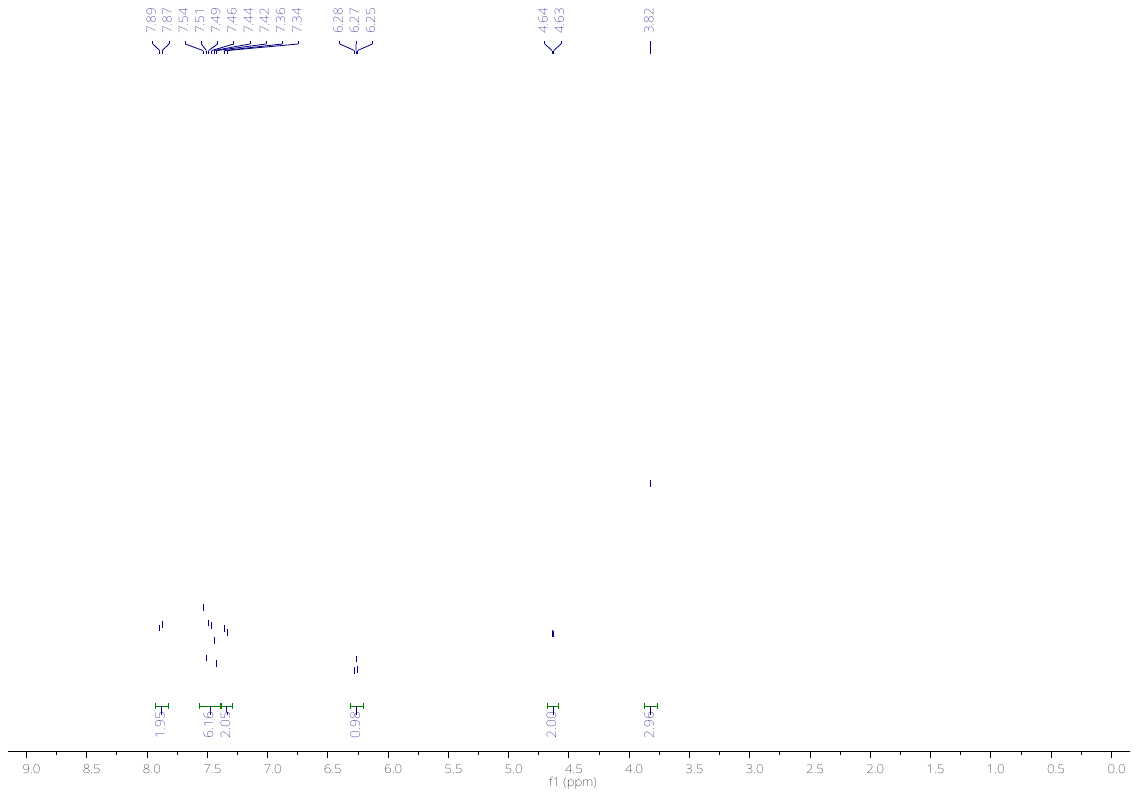

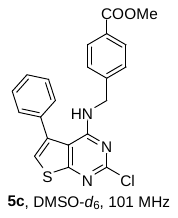

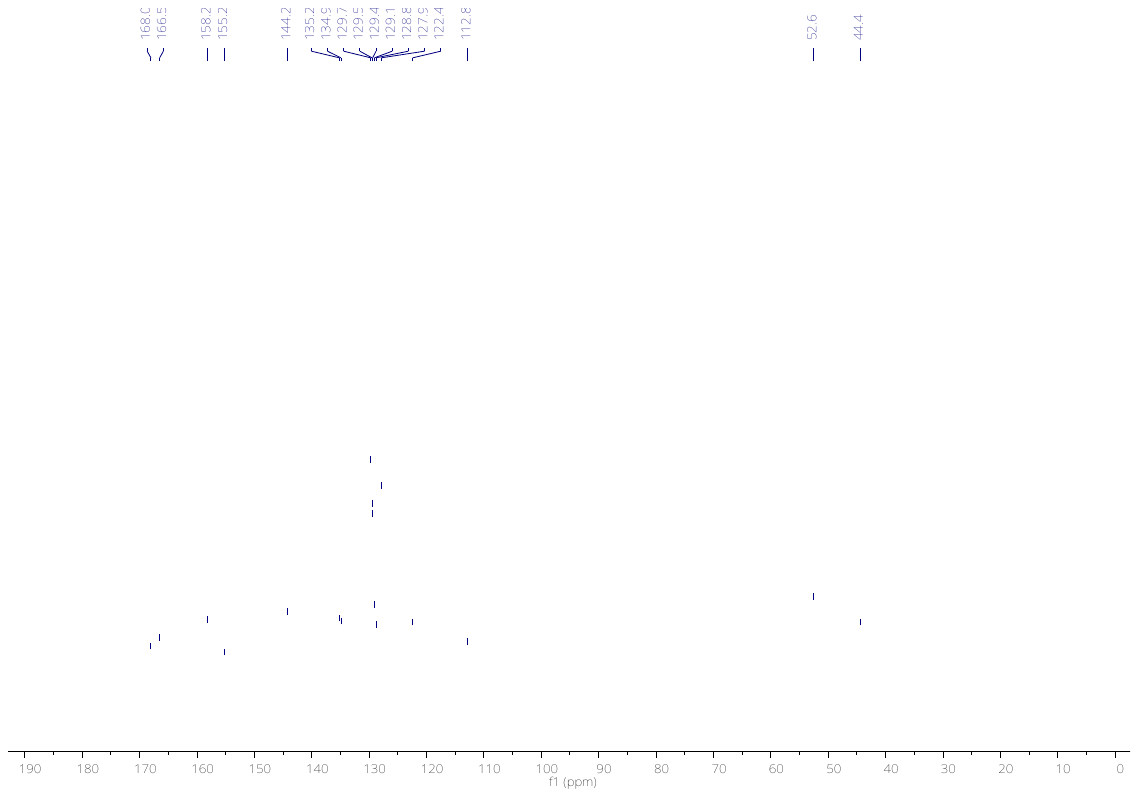

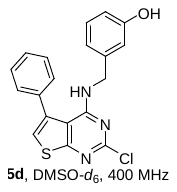

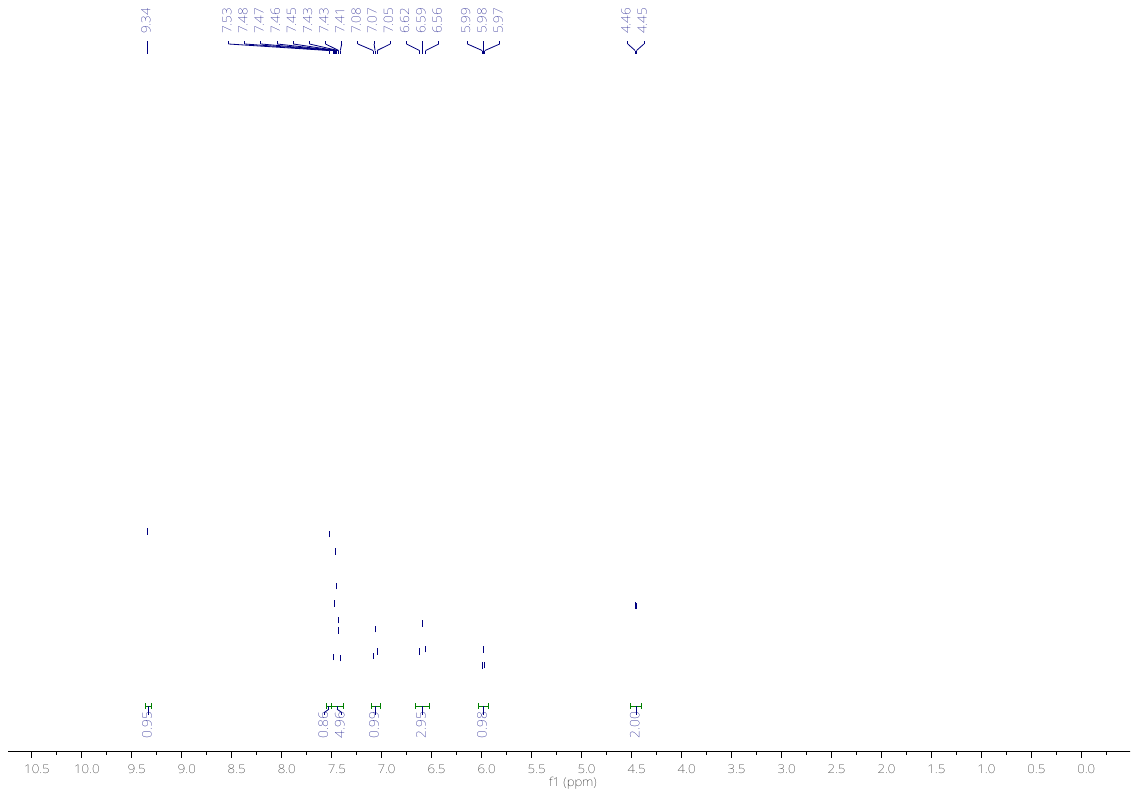

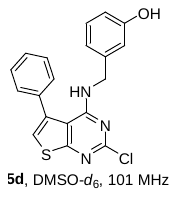

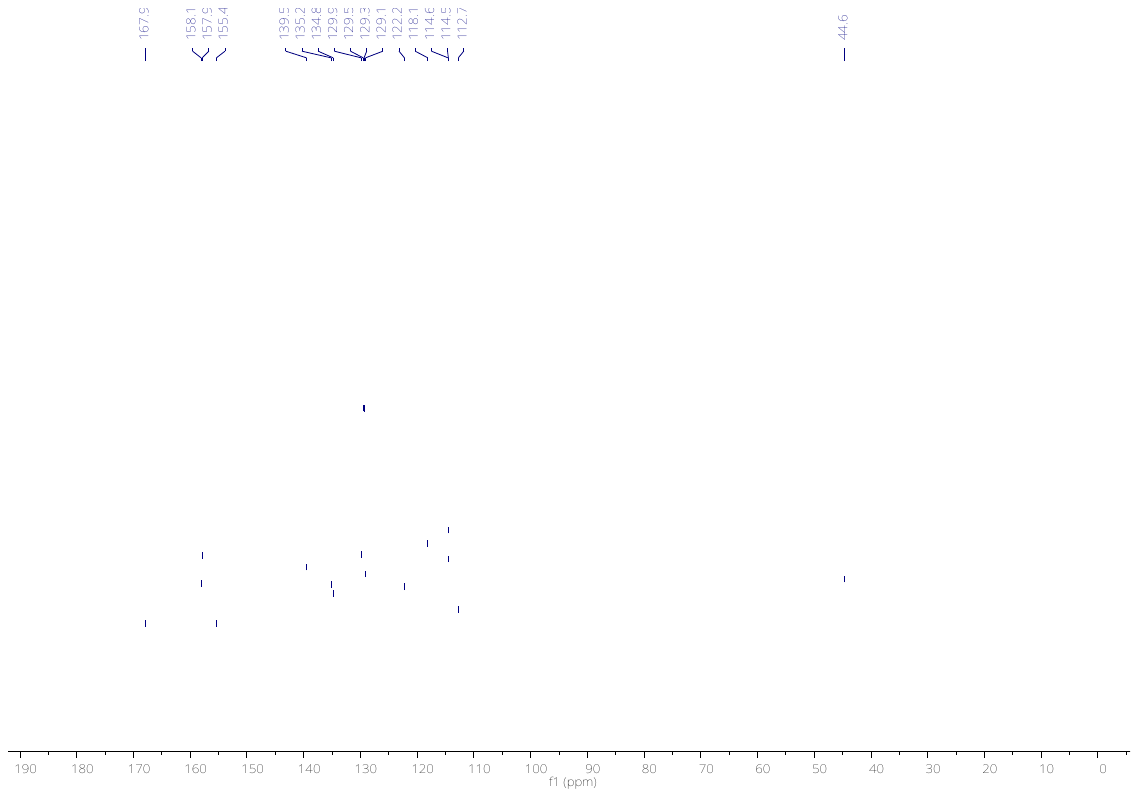

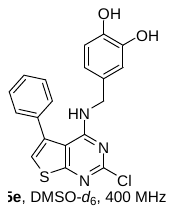

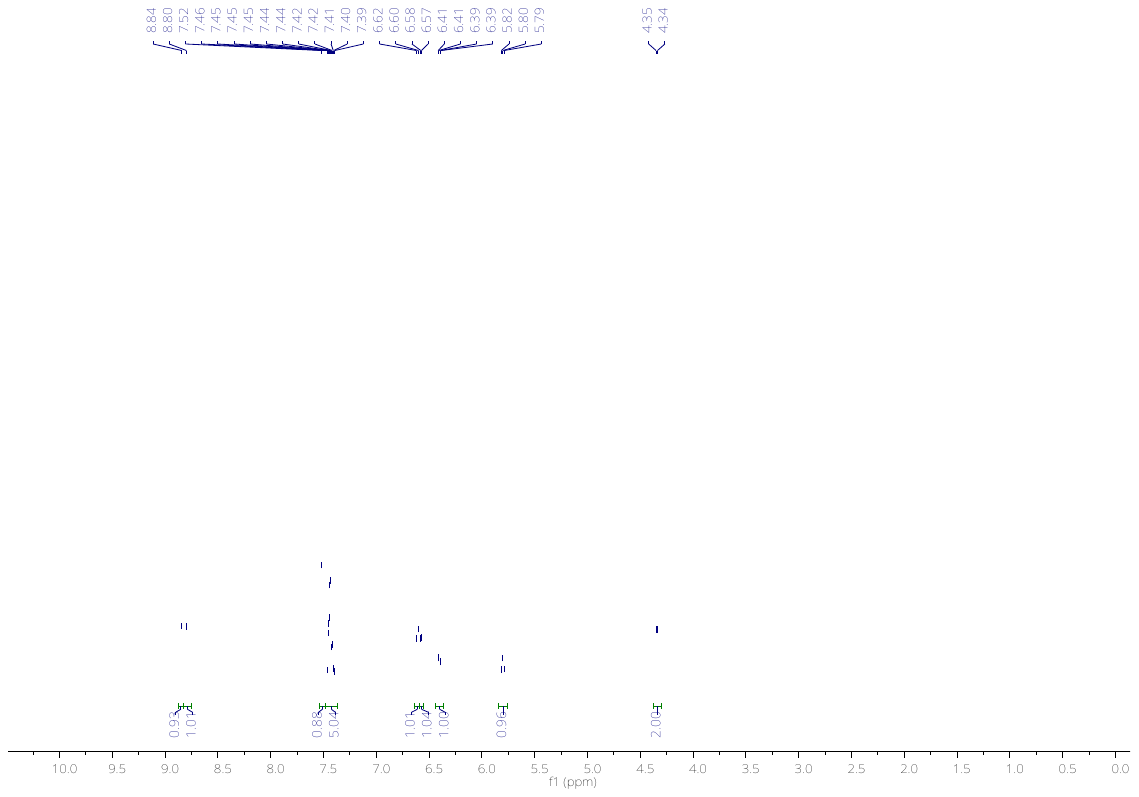

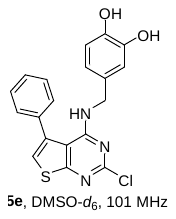

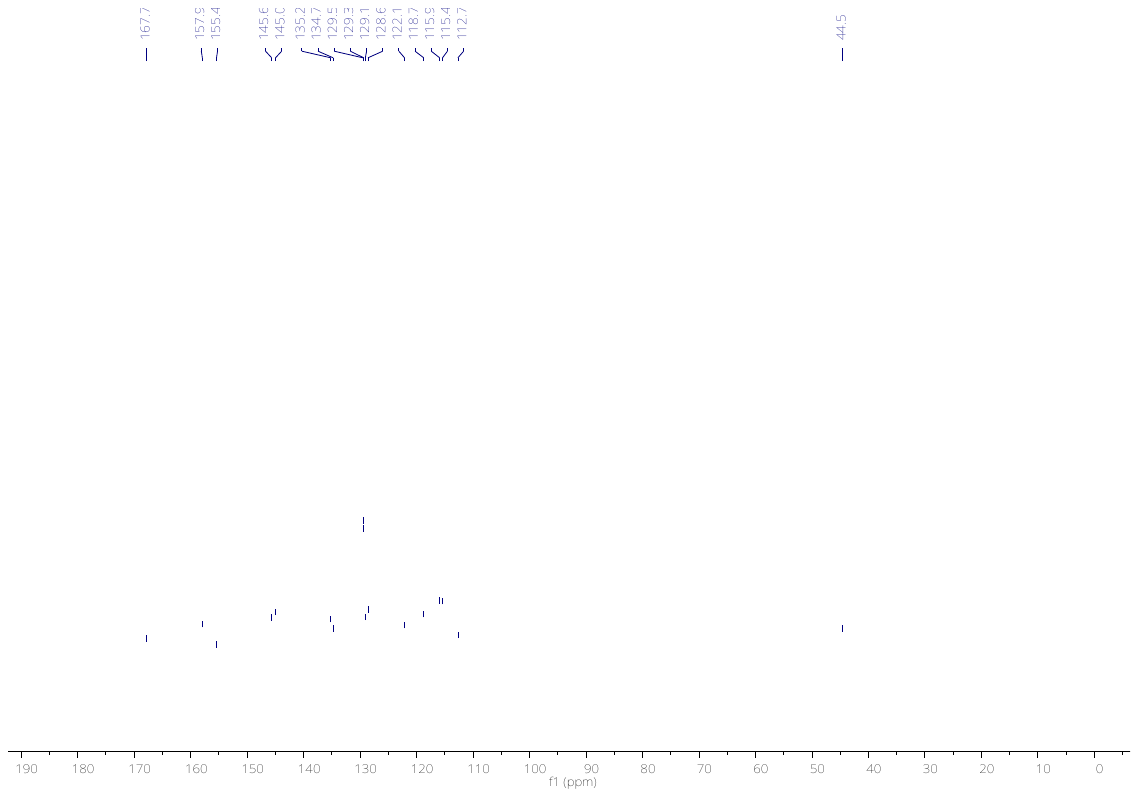

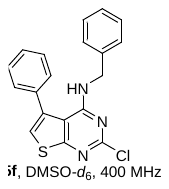

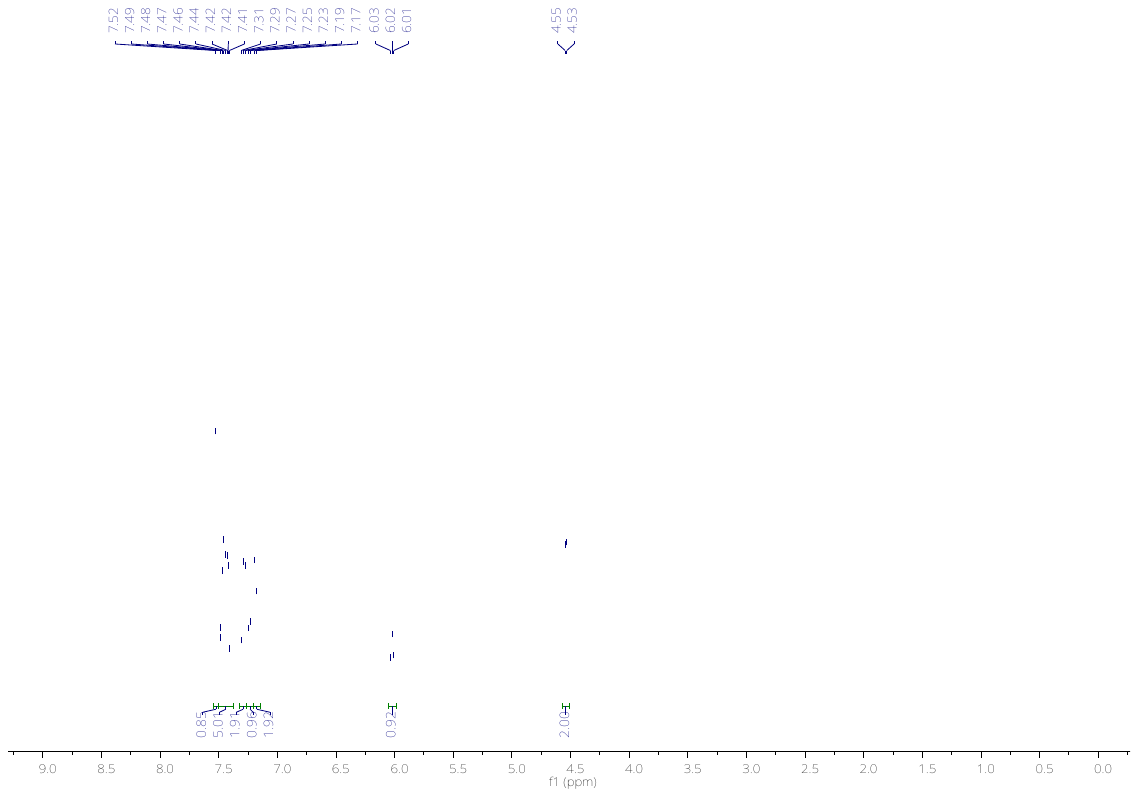

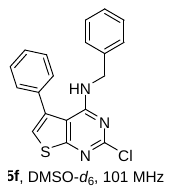

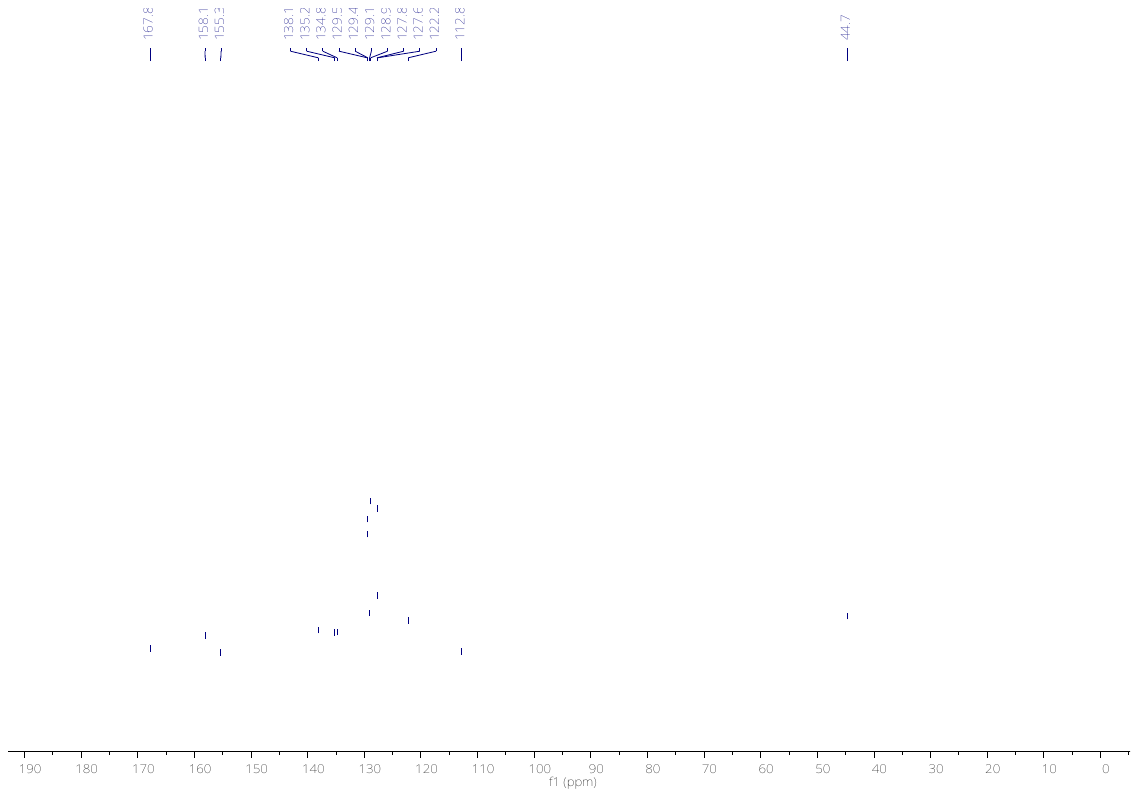

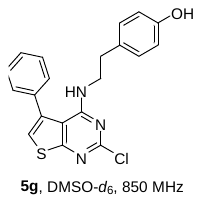

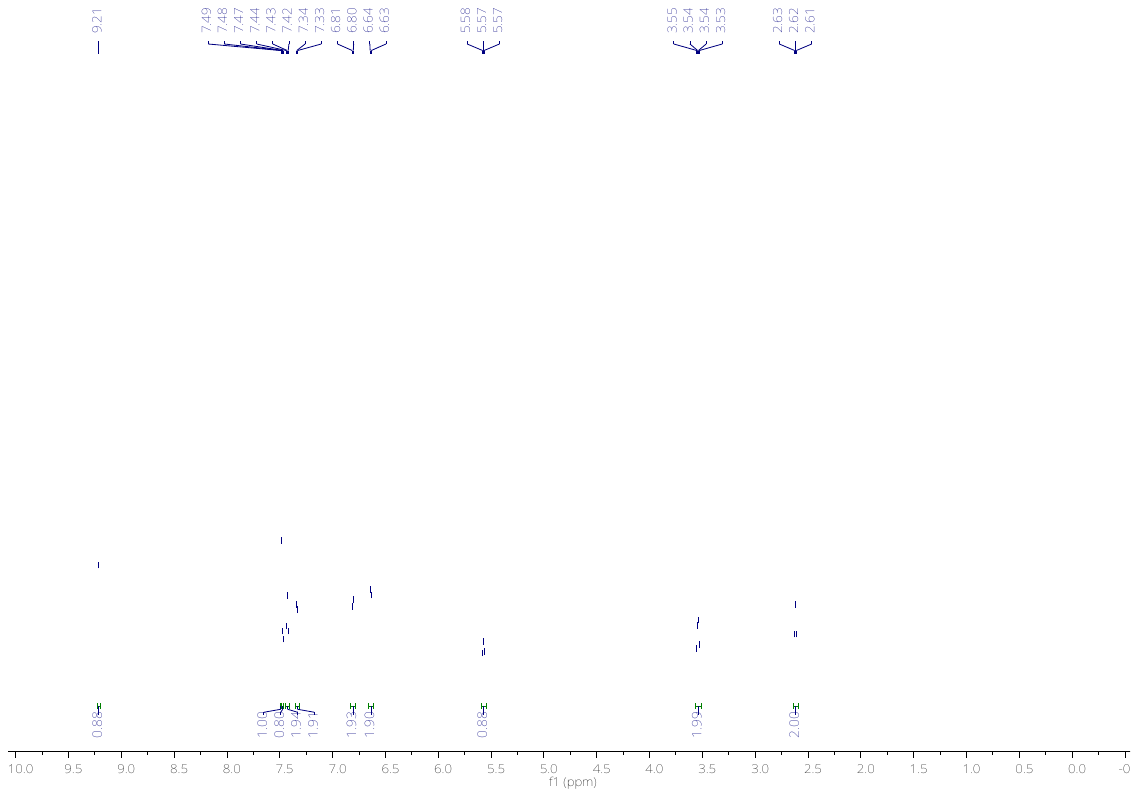

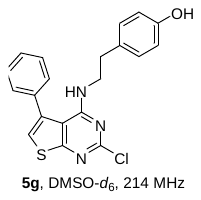

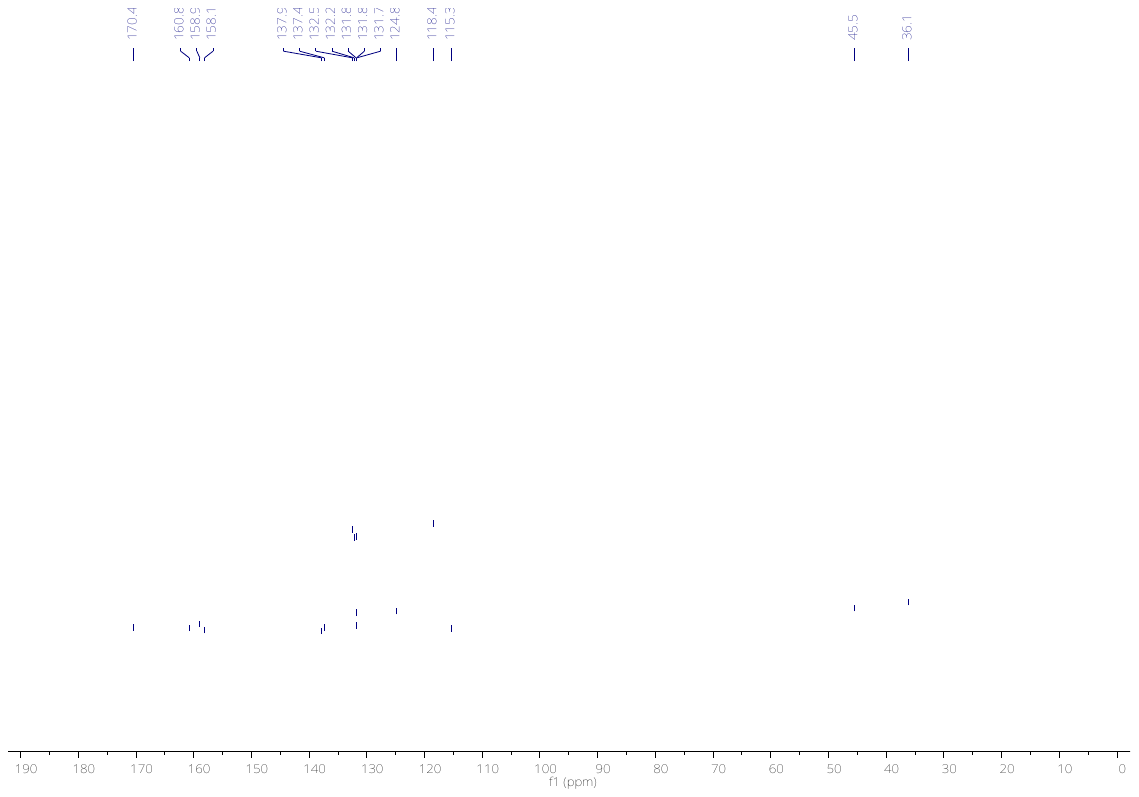

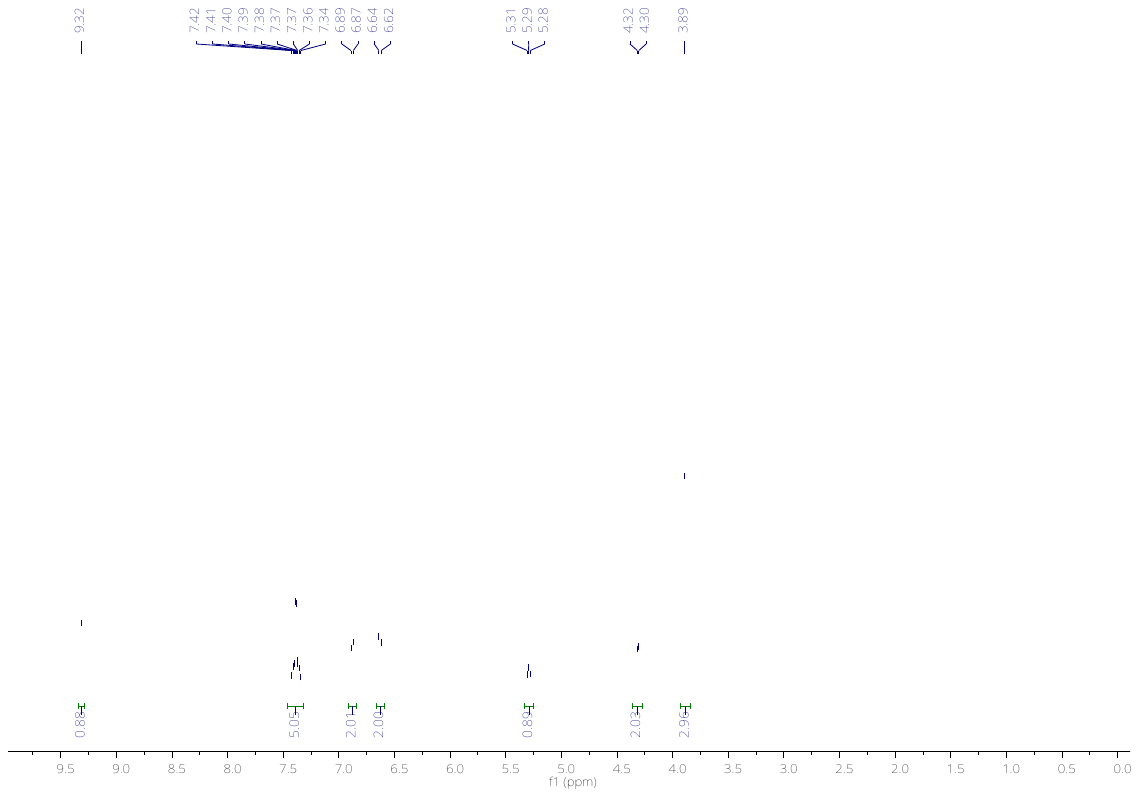

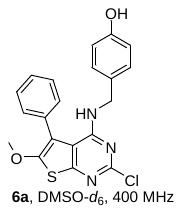

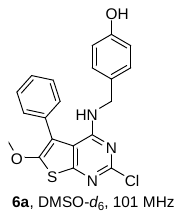

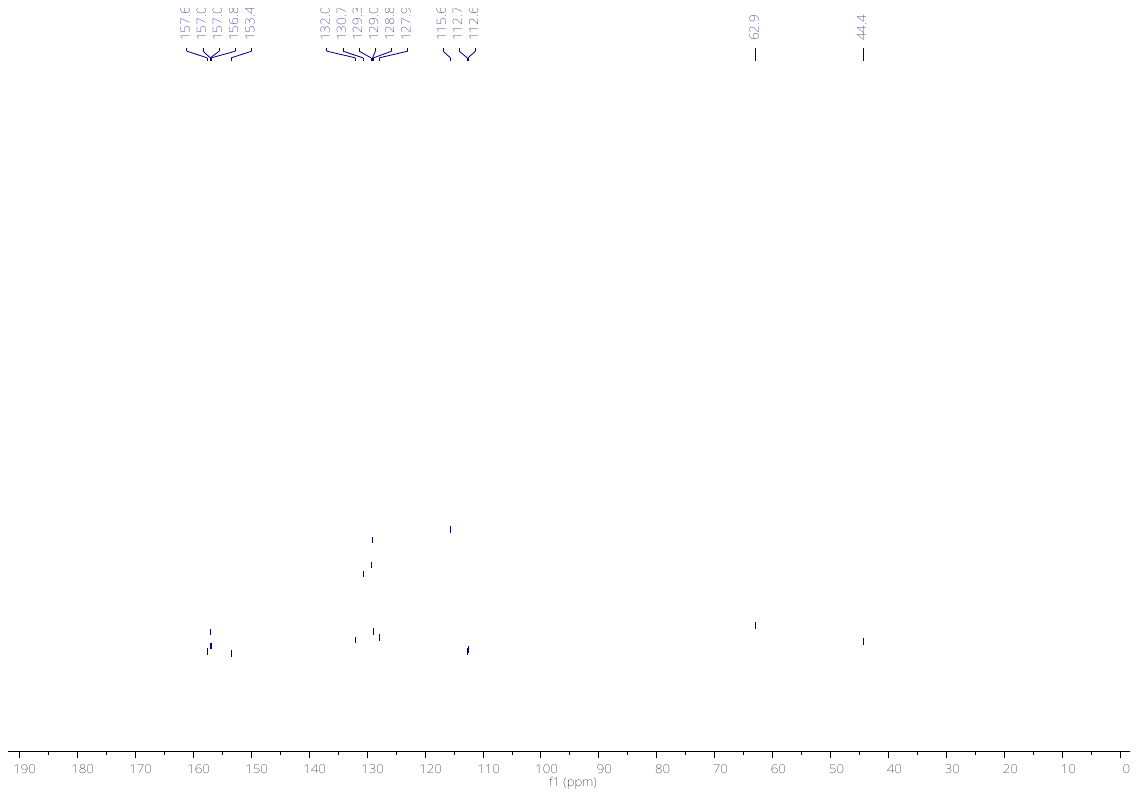

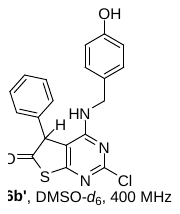

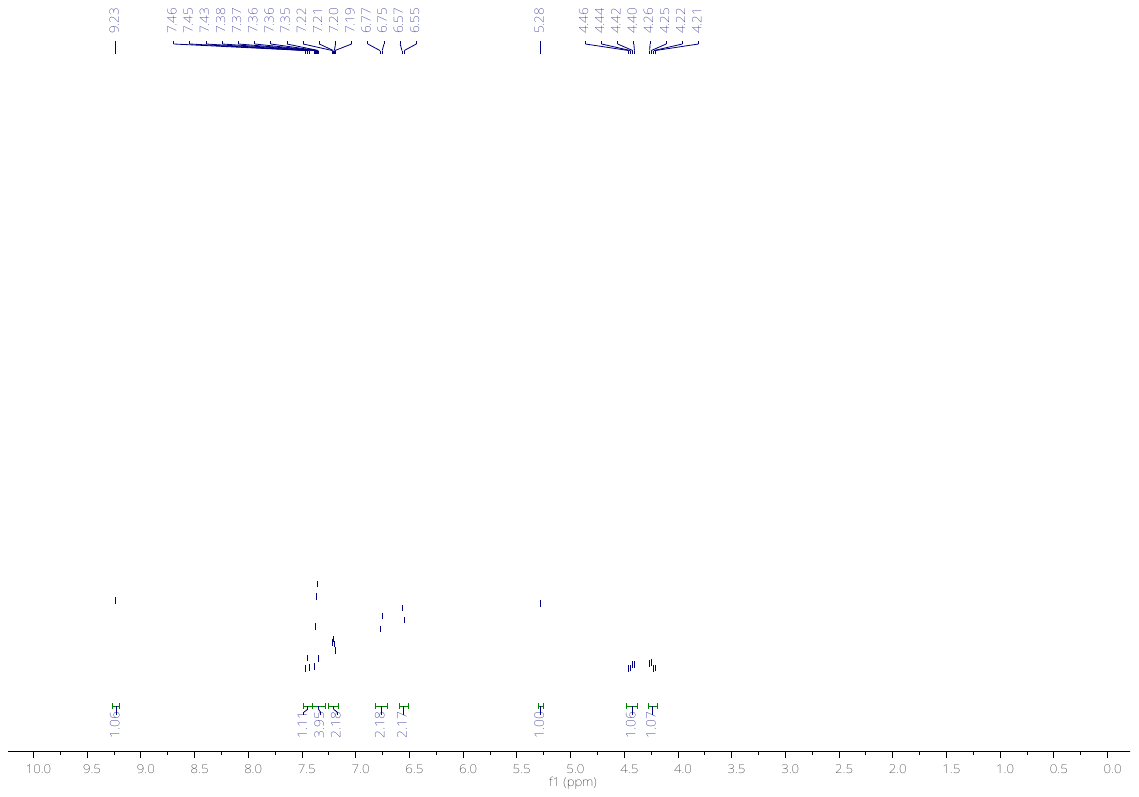

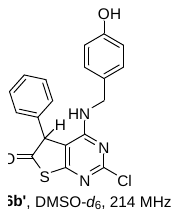

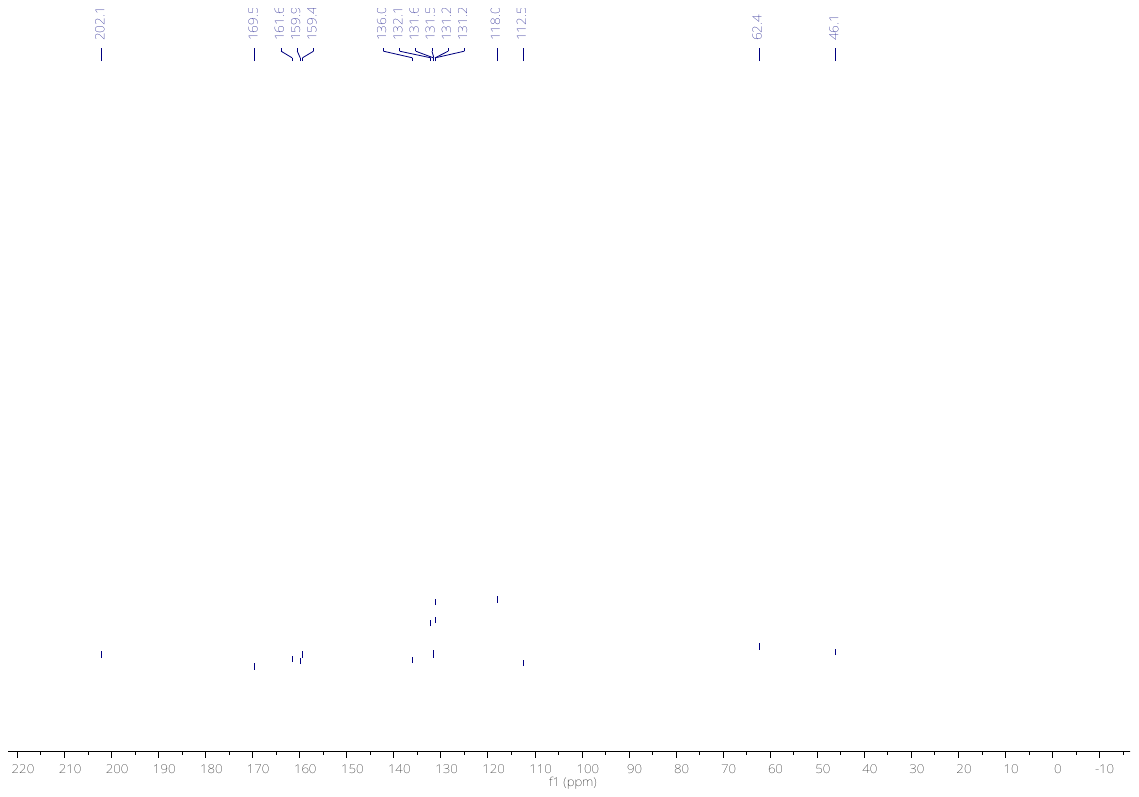
